# Functional mapping of androgen receptor enhancer activity

**DOI:** 10.1101/2020.08.18.255232

**Authors:** Chia-Chi Flora Huang, Shreyas Lingadahalli, Tunc Morova, Dogancan Ozturan, Eugene Hu, Ivan Pak Lok Yu, Simon Linder, Marlous Hoogstraat, Suzan Stelloo, Funda Sar, Henk van der Poel, Umut Berkay Altintas, Mohammadali Saffarzadeh, Stephane Le Bihan, Brian McConeghy, Bengul Gokbayrak, Felix Y. Feng, Martin E. Gleave, Andries M. Bergman, Colin Collins, Faraz Hach, Wilbert Zwart, Eldon Emberly, Nathan A. Lack

## Abstract

Androgen receptor (AR) is critical to the initiation, growth and progression of almost all prostate cancers. Once activated, the AR binds to *cis*-regulatory enhancer elements on DNA that drive gene expression. Yet, there are 10-100x more binding sites than differentially expressed genes. It still remains unclear how individual sites contribute to AR-mediated transcription. While descriptive functional genomic approaches broadly correlate with enhancer activity, they do not provide the locus-specific resolution needed to delineate the underlying regulatory logic of AR-mediated transcription. Therefore, we functionally tested all commonly occuring clinical AR binding sites with Self-Transcribing Active Regulatory Regions sequencing (STARRseq) to generate the first map of intrinsic AR enhancer activity. This approach is not significantly affected by endogenous chromatin modifications and measures the potential enhancer activity at each *cis*-regulatory element. Interestingly we found that only 7% of AR binding sites displayed increased enhancer activity upon hormonal stimulation. Instead, the vast majority of AR binding sites were either inactive (81%) or constitutively active enhancers (11%). These annotations strongly correlated with enhancer-associated features in both cell line and clinical prostate cancer. With these validated annotations we next investigated the effect of each enhancer class on transcription and found that AR-driven inducible enhancers frequently interacted with promoters, forming central chromosomal loops critical for gene transcription. We demonstrated that these inducible enhancers act as regulatory hubs that increase contacts with both other AR binding sites and gene promoters. This functional map was used to identify a somatic mutation that significantly reduces the expression of a commonly mutated AR-regulated tumour suppressor. Together, our data reveal a complex interplay between different AR binding sites that work in a highly coordinated manner to drive gene transcription.

## Introduction

Androgen receptor (AR)-mediated transcription is the primary driver of prostate cancer (PCa) growth and proliferation^1^. Activation of this critical signalling pathway occurs when AR binds to androgens such as testosterone or dihydrotestosterone (DHT). This induces the translocation of the AR into the nucleus, where it interacts with DNA at AR binding sites (ARBS). Almost all of these *cis*-regulatory elements (CRE) are located at distal intergenic or intronic regions^2, 3^. The AR cistrome is influenced by various transcription factors and pioneer factors, including FOXA1, HOXB13, and GATA2^2, 4, 5^. Once bound to DNA, the AR recruit numerous co-activators (CBP/p300, SRC/p160), chromatin modifiers (SWI/SNF-BRG1) and co-repressors (HDAC, NCoR) in a highly coordinated manner^6^. This protein complex then physically interacts with gene promoters via chromosomal loops, activating basal transcriptional machinery to drive transcription. Yet similar to other nuclear receptors, most AR-regulated genes interact with multiple ARBS^7^. There are vastly more ARBS (tens of thousands) than AR regulated genes (hundreds)^8, 9^. We do not know if these ARBS are inactive or active enhancers that interact in an additive, synergistic or dominant mechanism to induce gene transcription. Characterization of ARBS enhancer activity is critical to interpret the underlying regulatory logic of this transcription factor.

Enhancers have traditionally been identified by correlating transcription factor binding sites with chromatin accessibility, RNA polymerase II, GRO-seq or enhancer-associated histone modifications such as H3K27ac^10–13^. These features all broadly correlate with active enhancers but they are not causative and are therefore extremely prone to false positives^14^. For example, global loss of enhancer mark H3K27ac has no functional impact on gene transcription, chromatin accessibility or histone modifications^15^. Therefore, episomal reporter assays, which quantify the enhancer induced transcription of a gene, still remain the cornerstone of enhancer validation^16^. These assays are not influenced by endogenous chromatin compaction or epigenetic modifications and test the potential enhancer capability of each specific cis-regulatory element ^17^. While robust, conventional approaches are very low-throughput. To overcome these limitations several massively parallel reporter assays (MPRA) have been developed including Self-Transcribing Active Regulatory Regions sequencing (STARRseq)^18, 19^. In this approach, enhancer activity is quantified by measuring the rate of self-transcription of the genomic region cloned downstream of a minimal promoter. By utilizing multiplexed barcodes, the enhancer activity of many thousands of genomic regions can be measured simultaneously and provide locus-specific resolution of enhancer activity.

There is increasing clinical evidence that non-coding mutations can act as oncogenic drivers in PCa^20–22^. Recent studies by our lab and others have shown that ARBS are highly mutated in a tissue specific manner^23, 24^. Given the critical role of AR in PCa progression and treatment resistance any changes to the transcriptional landscape could alter tumour cell proliferation and sensitivity to AR pathway inhibitors. However, establishing a causal link between these mutations to a phenotype is extremely challenging due to the lack of functional CRE annotation, and the vast majority of these non-coding mutations remain unexplored in PCa. Better characterization of these CRE in PCa is essential to stratify potential driver mutations.

To provide the first locus-specific AR regulatory map we functionally quantified the enhancer activity of all commonly observed clinical ARBS with STARRseq. We demonstrate that only 7% of ARBS have androgen-dependent enhancer activation, while 11% have enhancer activity independent of AR binding. Surprisingly the vast majority of ARBS (81%) do not have significant androgen-dependent or constitutively active enhancer activity. These in vitro annotations strongly correlated to clinical PCa samples. To characterize the mechanism of AR enhancers we trained a machine learning classifier that can successfully predict active enhancers and identify key features for active enhancers. Integrating long-range chromatin interactome and transcriptomic data, we found androgen inducible enhancers were significantly more enriched as anchors for gene looping and acted as ‘hubs’ to activate AR-regulated genes. Finally, combining these results with whole genome sequencing of primary and metastatic PCa, we identified and characterized a non-coding somatic mutation that significantly impacted AR enhancer activity of a critical tumour suppressor.

## Results

### Functional quantification of AR enhancer activity

To functionally characterize AR CRE we experimentally tested the enhancer activity of all commonly occurring clinical ARBS sites with STARRseq, a massive parallel enhancer assay (**Figure 1A**). During optimization we found that ARBS inserts less than 250bp had marginal activity suggesting that the flanking sequences contribute to AR enhancer activity (**Supplementary Figure 3**). As the current synthesis limit of pooled oligos is ∼200bp we therefore used a capture-based approach to maintain a large insert size in the library. We designed a custom DNA capture assay to enrich the following genomic regions: common ARBS that are found in all normal tissue or primary PCa clinical AR ChIPseq^2^ (clinical ARBS; n=4139), previously identified strong enhancers that are not associated with AR^48^ (positive control; n=500) and regions where the AR does not bind in either clinical samples or cell lines yet contains an ARE-motif (negative control; n=2783). With this we then captured fragmented normal genomic DNA and cloned into a second-generation STARRseq plasmid^25^. A total of 365,265 unique on-target inserts (median 50 inserts/region) were cloned, with a normal distribution of inserts across our capture regions and a median insert size >500bp (**Supplementary Figure 4A**). Using this targeted library we tested for AR enhancer activity in an androgen-dependent PCa cell line (LNCaP). The resulting data demonstrated excellent reproducibility across biological replicas (*Pearson correlation 0.84-0.99*; **Supplementary Figure 4B**) and a strong STARRseq signal at known AR enhancers that regulate *KLK3* (**Figure 1B**). Similar to previously published work, our results for the STARRseq enhancer activity was comparable to a conventional luciferase reporter assay^18, 25^ (**Figure 1C**). Importantly, as STARRseq is a plasmid-based approach the enhancer activity is independent of endogenous chromatin compaction and quantifies the potential activity at each genomic region. This is clearly demonstrated with the non-AR positive controls which had strong enhancer activity regardless of being found in either heterochromatin or euchromatin^48^ (**Supplementary Figure 5**). When comparing all genomic regions tested, AR-driven enhancer activity was almost exclusively limited to ARBS with only 2/2783 ARE regions showing a significant increase in signal following androgen treatment **(Figure 1D**). Interestingly, within the ARBS regions we observed three distinct classes of enhancer CRE: a “classical” AR enhancer that increases activity when treated with androgen (inducible), enhancers that were active regardless of androgen treatment (constitutive) and those ARBS that had no significant enhancer activity (inactive) (**Figure 1E**). Of these, inactive CRE were by far the most common (81.9%; 3388/4139) with no enhancer activity either before or after androgen activation. A total of 11.2% (465/4139) and 6.9% (286/4139) were constitutively active or inducible enhancers, respectively. Surprisingly, none of the ARBS that were found only in clinical tissue but not LNCaP cells were inducible enhancers (n=867; **Supplementary Figure 6A+B**). This rate of inactivity is significantly less than expected compared to the clinical ARBS that overlap with LNCaP (*p<2.1×10^-16^*). As these regions lacked AR binding they were separated in subsequent analysis (no AR).

**Figure 1:**
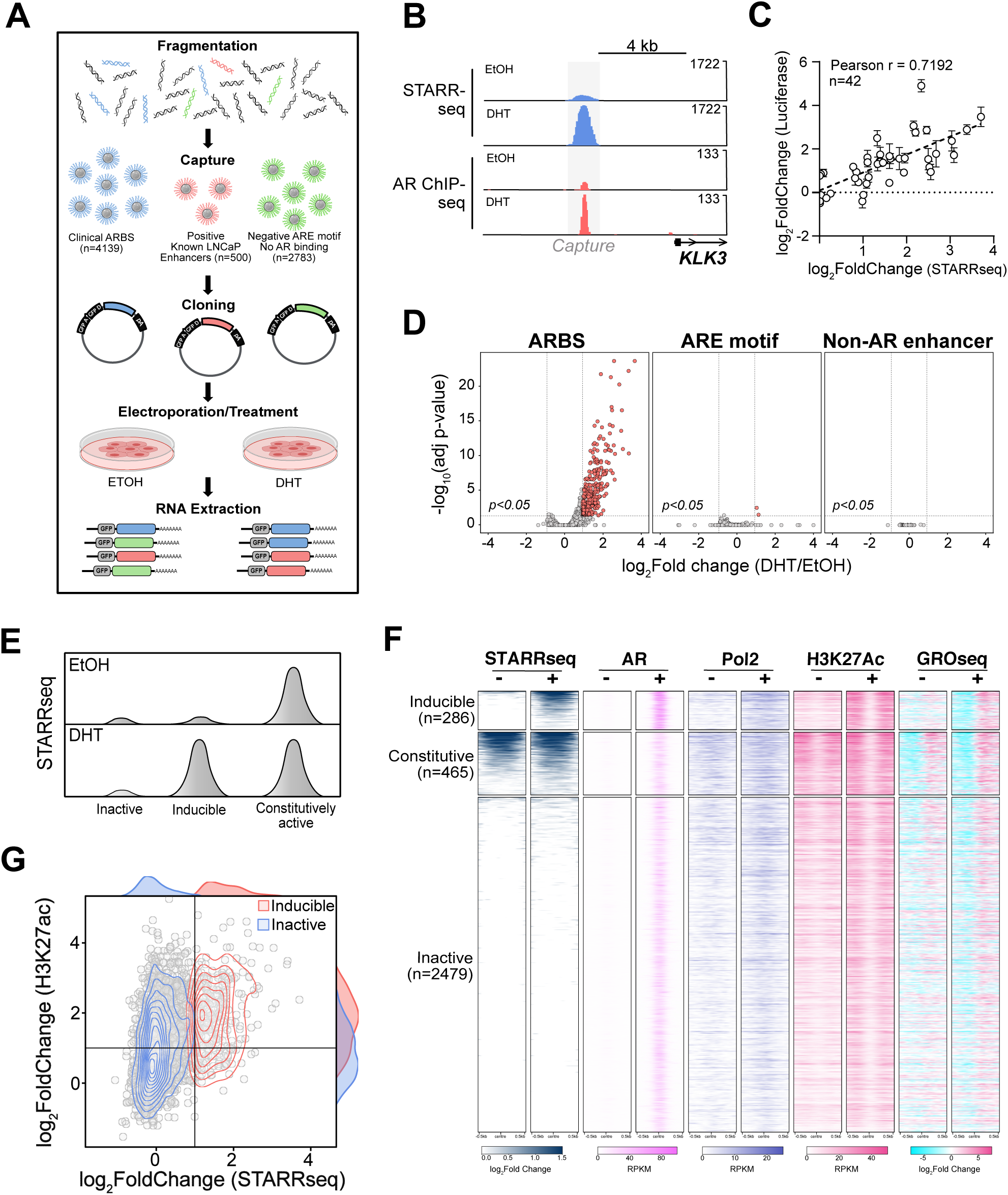
STARRseq identifies AR-dependent enhancers: **(A)** Schematic representation of AR STARRseq. **(B)** Strong androgen-dependent enhancer activity (red) was observed at known AR binding sites (blue; GSE83860) proximal to *KLK3*. **(C)** Enhancer activity of AR CRE with varying levels of STARRseq signal (n=42) were validated with a luciferase assay (4 biological replicates±SEM). A strong correlation is observed between luciferase and STARRseq signals. **(D)** Volcano plot of androgen-dependent changes in STARRseq enhancer activity for clinical ARBS (n=4139), ARE motif alone (n=2783) and strong non-AR enhancers (n=500). Significantly induced enhancers (LFC≥1, *p-adj<0.05*) are highlighted in red. **(E)** Schematic representation of the different classes of AR enhancers. **(F)** Heatmap of STARRseq, publicly available ChIPseq of AR, Pol2, and H3K27ac as well as GROseq in EtOH or DHT treated LNCaP cells. The heatmap is divided based on the functional classes of each enhancer class identified by STARRseq. **(G)** Density map of androgen induced changes to H3K27ac ChIPseq and STARRseq at inactive and inducible AR enhancers.

To confirm that our AR CRE annotations correlated with enhancer activity *in vitro* we compared each group with published enhancer-associated features including H3K27ac, RNA polymerase II (Pol2) and bidirectional eRNA (GROseq) (**Figure 1F**). We observed a strong correlation between each enhancer group and these features. Specifically, constitutive AR enhancers demonstrated high levels of H3K27ac, Pol2 and eRNA that were comparable between androgen -deprived and androgen containing conditions. In contrast, for inducible enhancers these features only increased when cells were treated with androgens. For inactive CRE, enhancer associated features were broadly reduced compared to active enhancers though there was some variation observed. Further, we found that there were marked differences in AR-mediated DNase I hypersensitive sites between different AR CREs, with inducible enhancers dramatically increasing accessibility following androgen treatment (**Supplementary Figure 7A**). There was no significant enrichment for any one AR enhancer class at super enhancers as these elements are relatively rare at ARBS (**Supplementary Figure 7B**). Yet while these descriptive features generally correlate with active AR enhancers, they are extremely prone to false positives at individual CRE. For example, while inducible AR enhancers generally have higher androgen-induced H3K27ac than inactive ARBS there is significant overlap between these classifications (**Figure 1G**). Given that inactive CREs are far more common than induced enhancers this dramatically increases the false positive rate. Specifically, if an active enhancer is called solely on AR and H3K27ac ChIPseq there is a >80% false positive rate. Supporting these results, the enhancer activity of high H3K27ac inactive and inducible CRE was validated with a luciferase reporter assay (**Supplementary Figure 8**). Overall these results demonstrate that functional enhancer testing is needed to provide locus-specific resolution and annotate CRE.

### Clinical validation of enhancer annotation

While histone modifications do not accurately identify individual AR enhancer CREs (**Figure 1G**), these features, particularly H3K27ac, do broadly correlate with active enhancers (**Figure 1E**). Therefore to determine if our enhancer annotations represented clinical AR activity, we analyzed previously published AR (n=87), H3K27ac (n=92) and H3K27me3 (n=76) ChIPseq from primary PCa tissue^8^. Supporting our *in vitro* classifications we observed significant enrichment of both AR and H3K27ac at inducible and constitutive enhancers as compared to inactive ARBS (**Figure 2A**). Further, while not as dramatic, we also found a statistically significant enrichment of the repressive H3K27me3 mark at inactive ARBS compared to constitutive and induced enhancers (*p-adj<0.05)*. However, as these primary PCa samples contain physiological levels of androgen we could not separate induced and constitutive enhancers. Therefore, to further validate our *in vitro* classifications we conducted H3K27ac ChIPseq on prostate tumours from patients enrolled in a neoadjuvant antiandrogen ENZA clinical trial (NCT03297385). Tumour samples were collected pre-and post-ENZA thereby allowing the impact of AR activity to be quantified in matched clinical samples. As expected, ChIPseq results from pre-ENZA patients were very similar to the primary PCa samples with an enrichment of H3K27ac in constitutive and inducible ARBS as compared to inactive ARBS (**Figure 2B**). However, following ENZA treatment H3K27ac was only enriched at constitutive enhancers and both induced and inactive CRE had markedly lower histone modifications (**Figure 2B**). When normalized to constitutive and inactive CRE, ENZA treatment markedly reduced H3K27ac at inducible AR enhancers (**Figure 2C)**. Overall these results demonstrate that our *in vitro* classifications strongly correlate to clinical AR activity and suggest that this plasmid-based enhancer assay represents AR activity *in situ*.

**Figure 2:**
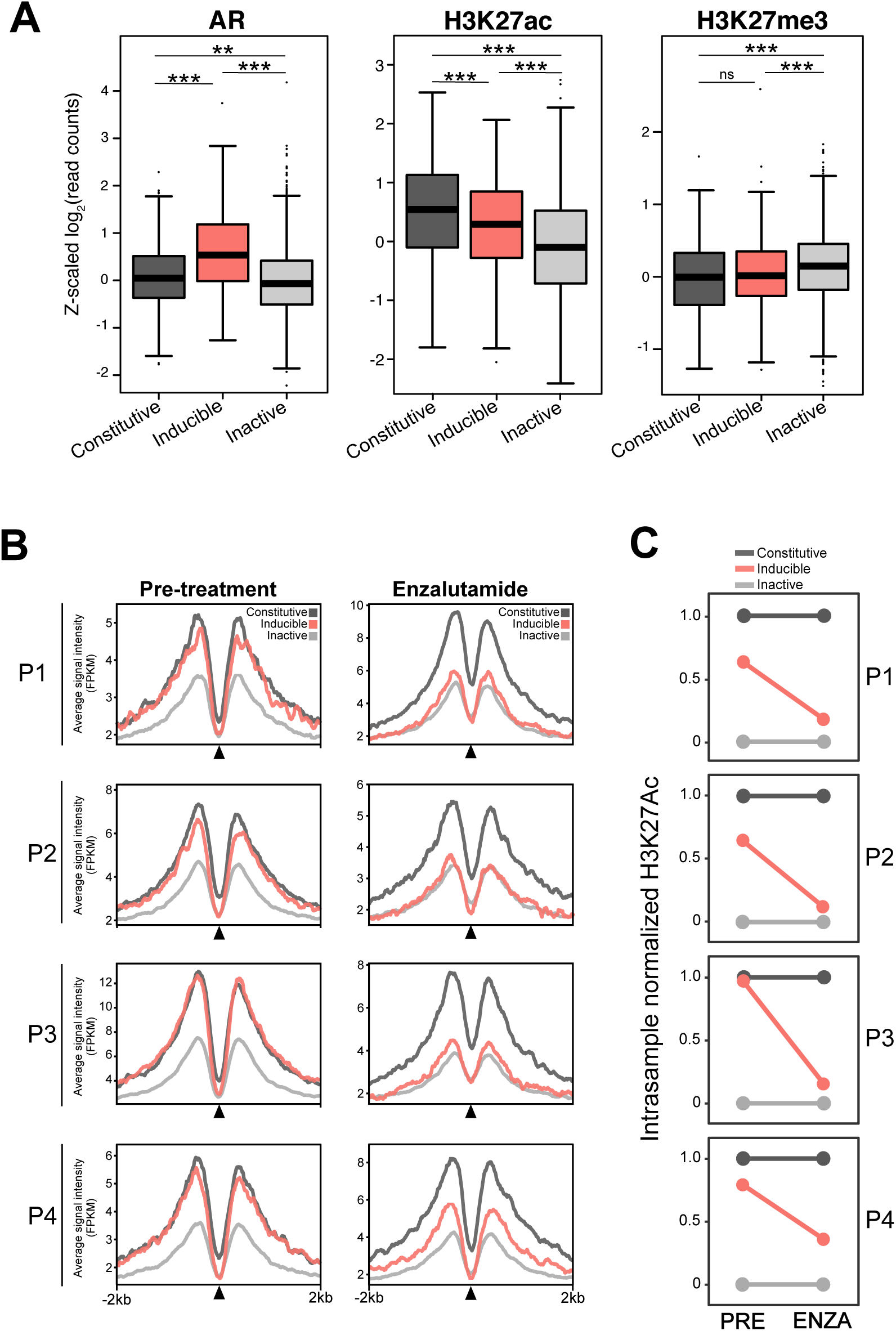
*In vitro* enhancers classification are preserved in clinical samples: **(A)** Normalized ChIPseq of AR (n=87), H3K27ac (n=92) and H3K27me3 (n=76) from primary PCa samples. A significant enrichment of AR and H3K27ac are observed in induced and constitutive compared to inactive enhancers (*ns>0.05, ** p<10^-4^, *** p<10^-6^*). H3K27me3 was also significantly enriched in inactive enhancers compared to induced or constitutive enhancers. **(B)** H3K27ac ChIPseq was done in matched patient PCa tissue pre- and post-enzalutamide treatment (n=4). Normalized H3K27ac enrichment ± 2Kb around induced (red) constitutive (dark grey) and inactive (grey) enhancers are shown. **(C)** H3K27ac enrichment in each class of enhancers were normalized within each tumour and the normalized scores were compared before and after ENZA treatment. H3K27ac enrichment in induced enhancers was markedly reduced after ENZA treatment.

### Genomic features associated with AR enhancers

Having mapped the AR CRE enhancer activity, we next analyzed the DNA motifs at each ARBS to determine what feature correlated with active enhancers. Unfortunately, this gave very poor results with almost no difference in DNA motifs including AREs at inactive, inducible and constitutively active ARBS (**Supplementary Figure 9A**). This matches with our experimental findings, where almost all genomic regions containing an ARE motif without AR binding had no inducible enhancer activity (**Figure 1D**). Expanding on our observation that AR binding in LNCaP was required for AR-mediated enhancer activity (**Supplementary Figure 6**), we incorporated all publically available experimental genome-wide ChIPseq data from LNCaP cells (**Supplementary Table 2;** n=90) and trained a machine learning classifier to predict enhancer activity at each ARBS (**Figure 3A**). All transcription factor and histone ChIPseq was processed and normalized with a standardized bioinformatic pipeline to reduce technical variation. Using this functional genomic information our bootstrapped multinomial logistic regression model with a sparsity LASSO regularizer achieved 65% precision. On test data, our model managed a 65% precision for the inducible group and a 62% precision for the inactive group with an overall accuracy of 60%. Given the low frequency of induced enhancers this is a >10x enrichment compared to random ARBS. To validate this predictive model we experimentally tested those LNCaP-specific ARBS not included in our clinical STARRseq library (**Figure 3B**). Confirming the test results, we observed that 60% of the predicted induced enhancers could be accurately identified with our classifier. As this uses a relatively simple multinomial logistic regression model we can quantify the predictive strength of each DNA-bound factor and identify those features that strongly correlate with inducible AR enhancers. When calculating the differential binding energy for the inducible and non-inducible groups we observed that most features associated with inducible enhancers were unsurprisingly found in androgen treated conditions. Specifically, AR, PIAS1, ARID1A, MED1, and RUNX1 binding peaks in androgen treated conditions were strong predictors of inducible AR enhancers (**Figure 3A**). In contrast, occupancy of CTBP2, WDHD1, and TLE3 at ARBS in EtOH were generally predictive of inactive CRE though these did not have comparable predictive strength to inducible features. To identify which functional genomic features best predicted inducible AR enhancers we downsampled our model by reducing the number of features and then re-tested each classifier compared to the general model. With this, we found that of all individual features, AR+DHT peak height had the best power to identify inducible enhancers at ARBS and was significantly better than either H3K27Ac or any other transcription factors/histone marks (**Figure 3C;** 0.81 vs. 0.55 AUC). If expanded to three features, ChIPseq of AR+/-DHT and PIAS1+DHT gave the best results with comparable recall to the larger general model (**Supplementary Figure 9B;** 0.83 vs. 0.86 AUC). Overall this machine learning classifier provides a powerful tool to both identify both those regions that are likely to be inducible enhancers and also the specific features associated with active AR enhancers.

**Figure 3:**
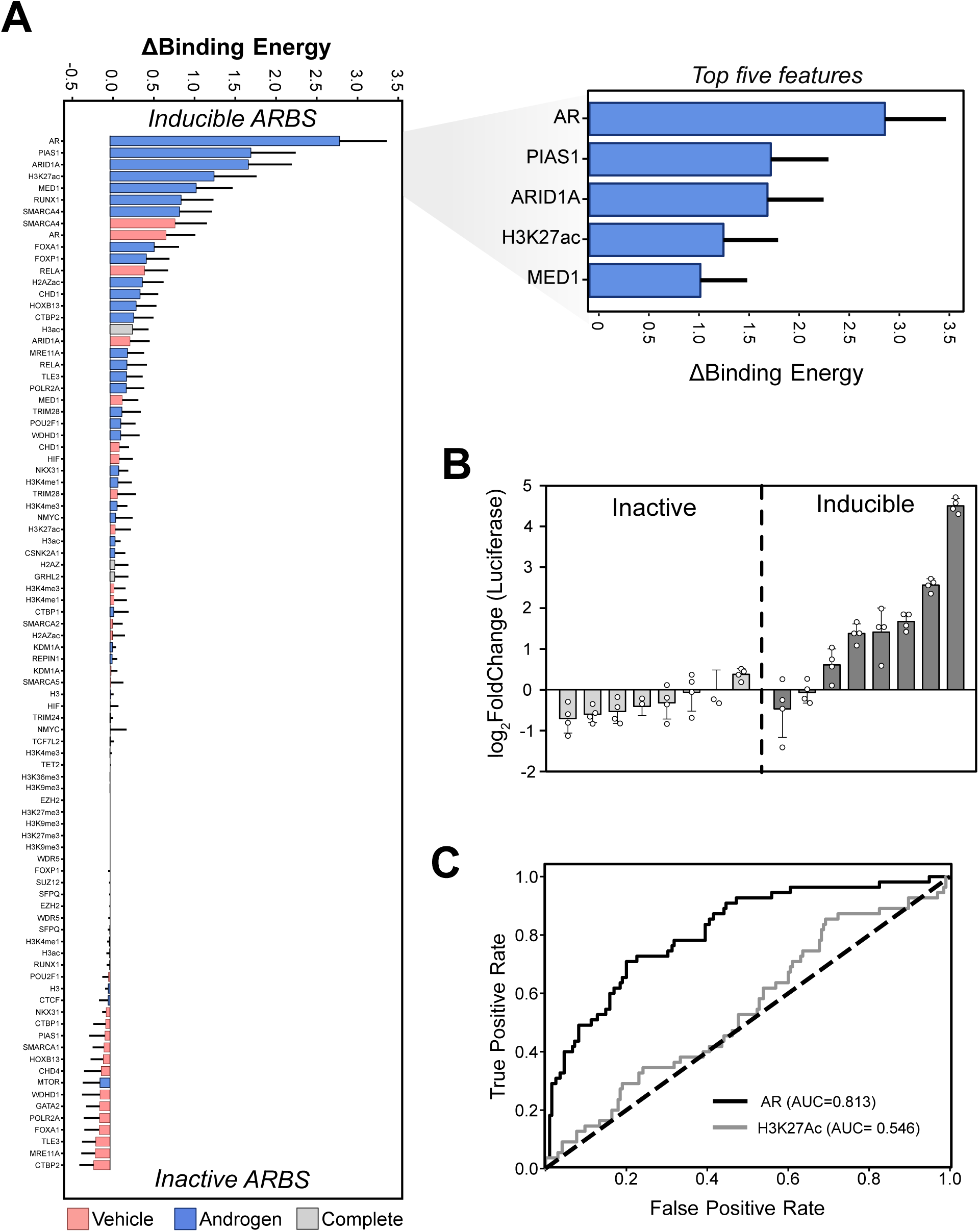
Identification of features associated with AR enhancers: **(A)** Features from all publicly available TFs and histone mark ChIPseq datasets were ranked based on their calculated binding energy at inducible enhancers. The zoomed section (top right) shows the top 5 features that are predictive of inducible enhancers. **(B)** LNCaP ARBS predicted by the classifier as either inactive or inducible enhancers were validated by luciferase assay. (4 biological replicates±SEM) **(C)** Receiver operating characteristic curve of AR and H3K27ac ChIPseq to accurately identify inducible enhancers

### Role of AR enhancers on gene expression

We next investigated how each AR enhancer class impacts androgen-mediated transcription. As enhancer-promoter (E-P) interactions frequently occur within neighbouring primary sequences^49, 50^, we first correlated the distance between promoters of androgen-induced genes to each AR enhancer class. We observed that inducible enhancers were significantly more likely to be near an androgen-upregulated differentially expressed genes (DEG) than a constitutive or inactive enhancer (**Figure 4A**). However, these E-P interactions frequently occur over considerable distances in the primary sequence due to chromatin looping^51^. Therefore, incorporating a published AR ChIA-PET dataset we characterized the chromatin loops between individual ARBS and AR regulated genes and found that inducible ARBS interact more frequently with promoters at androgen-upregulated DEG than either constitutive or inactive ARBS (*p<0.0001*; **Figure 4B**). No enrichment was observed between any enhancer class and androgen down-regulated DEG suggesting that this occurs through an indirect mechanism. We also observed that the inducible AR enhancers form significantly more chromatin loops to either other ARBS or chromosomal sites suggesting that they may act as regulatory “hubs” (*p<2×10^-^*^16^; **Figue 4C**). To better quantify the relationship between AR CREs, we transformed the pairwise interactions into an undirected network with all LNCaP ARBS or DEG TSS being represented as a vertex and the chromatin loops as edges (**Figure 4D**). All ARBS were including even those not tested by STARRseq to provide a more comprehensive AR interaction network. Matching our earlier analysis, we found that induced AR enhancers have a significantly enriched interaction frequency with upregulated DEG (**Figure 4E**). When relationships between AR CREs in connected or independent networks were quantified, we found that inducible enhancers were significantly more likely to be a central node in this regulatory network (*p<2×10^-9^;* **Figure 4F, Supplementary Figure 10A**). A similar trend was observed when comparing networks of only tested ARBS regions (**Supplementary Figure 10B**). These findings suggest that inducible enhancers may play a critical role in AR-mediated transcription as a central node between promoters and AR CRE.

**Figure 4:**
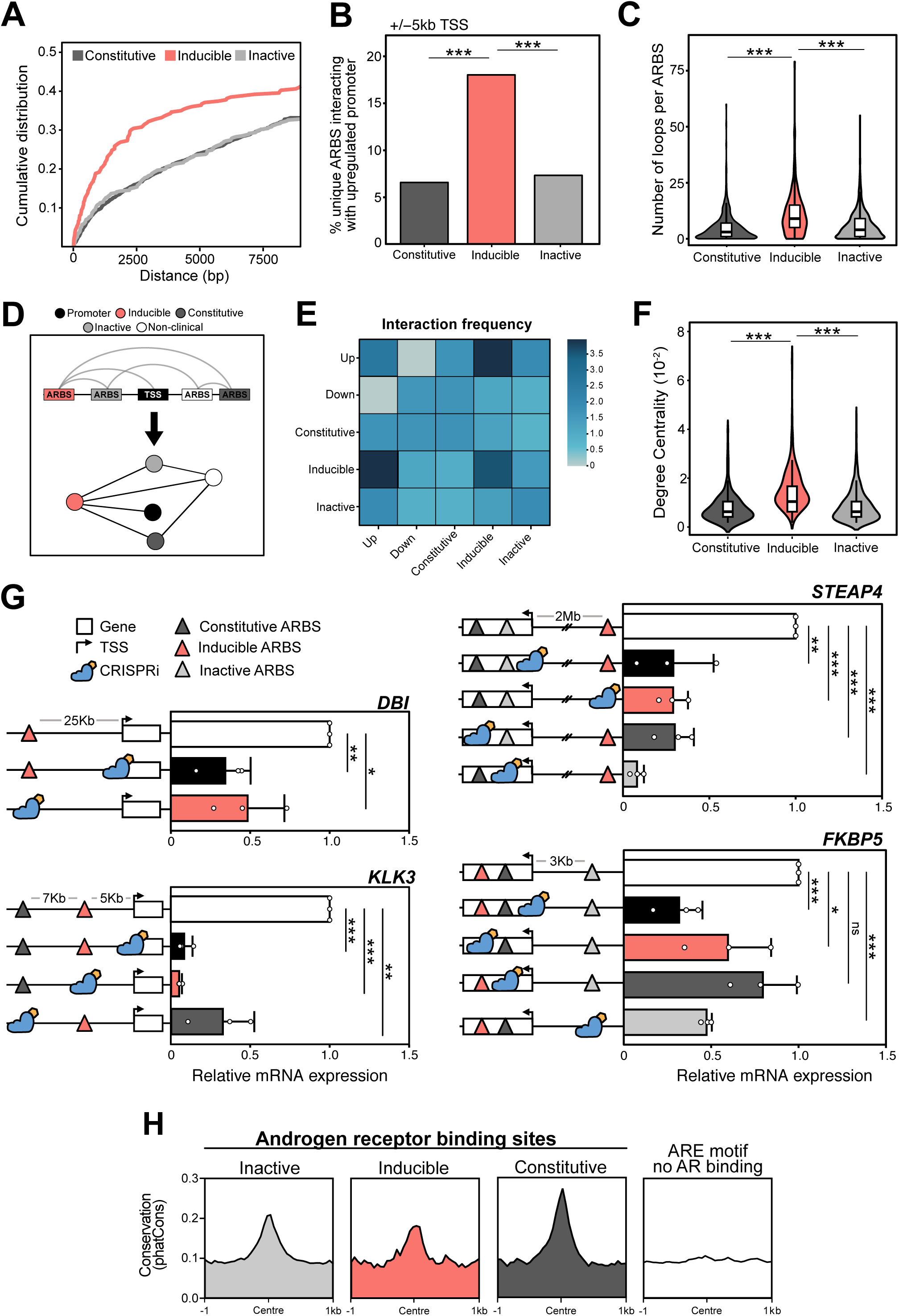
Transcriptional regulation by AR enhancers: **(A)** Cumulative distribution function correlating the distance (bp) of each enhancer class to the promoters of androgen upregulated genes. **(B)** Chromatin loops at ARBS (ChIA-PET) were overlapped with the enhancer classification to identify those ARBS that looped to a promoter (± 5kb from the TSS) of an androgen upregulated gene **(C)** The number of AR chromatin interactions from each enhancer class to any site was quantified for each enhancer class. **(D)** Schematic representation of AR ChIA-PET data transformed into graph network. **(E)** Calculation of the relative interaction frequency in the graph network between androgen up regulated gene promoters (Up), androgen down-regulated gene promoters (Down) and each ARBS enhancer class. (**F**) With the interaction graph network the betweenness centrality in the largest connected graph was calculated for each enhancer class (*ns p>0.05, *** p<10^-9^*). (**G**) At AR regulated genes, individual ARBS CRE (induced, inactive, constitutive) were inhibited with CRISPRi in LNCaP cells to determine their impact on androgen-mediated transcription. The changes in the gene expression were quantified by the qPCR. (3 biological replicates±SD;*\*\*\* p<10^-9^*). (**H**) Evolutionary conservation from 100 vertebrate species of different ARBS enhancer classes compared to ARE-motif alone genomic regions (n=2783).

To functionally test these descriptive results, we used CRISPRi to selectively inhibit annotated CRE and quantify the impact on AR-mediated gene expression^52^. Genes were chosen based on their enhancer complexity ranging from a single inducible enhancer that loops to the gene promoter (*DBI*) to a mixture of inactive, constitutive and inducible AR enhancers (*KLK3, STEAP4, FKBP5)* (**Figure 4G; Supplementary Figure 11A)**. Inactivation of inducible AR enhancers significantly reduced androgen-mediated transcription of all genes tested (**Figure 4G**). Similar to previously published work, when the *KLK3* inducible AR enhancer (AREIII) was inactivated it also decreased the neighbouring gene *KLK2*^53^. Highlighting the specificity of this approach, inactivation of the *KLK3* promoter did not downregulate *KLK2* (**Supplementary Figure 11B**). Surprisingly, in addition to the inducible enhancers, many of the constitutive and inactive ARBS also contributed to AR-mediated gene transcription. Inhibition of the *KLK3* constitutive enhancer significantly impaired gene expression suggesting a co-operative action between these two enhancers. In contrast, inhibition of a constitutive enhancer did not significantly alter androgen-mediated transcription of *FKBP5* while targeting either the AR inducible enhancer or inactive ARBS reduced expression. Finally, AR-mediated transcription of *STEAP4* was significantly reduced by inhibiting inactive, constitutive or inducible CRE. This is particularly striking as the *STEAP4* inducible AR enhancer forms a chromosomal loop to this TSS over a >2Mb distance (**Supplementary Figure 11A**). This functional testing suggests that while inducible enhancers are critical for gene expression, other AR enhancer classes can also be required to drive transcription. Supporting this potential role, inducible, inactive and constitutive CRE are all evolutionarily conserved compared to random ARE-motif regions (**Figure 4H**). This is unlikely to be an artifact of the STARRseq assay as the enhancer annotation classes strongly correlate with enhancer features both *in vitro* and *in vivo*.

Given these results, we proposed that inducible enhancers may act as a regulatory hub between multiple ARBS and gene promoters. To test this we conducted single cell ATACseq (scATACseq) to identify co-accessible AR CREs and determine the *cis*-regulatory interactions (**Supplementary Figure 12**). Co-accessible DNA elements strongly correlate with physical proximity^44^ and can characterize how AR binding changes CRE interactions, something that would not be possible with AR HiChIP or ChIA-PET. Similar to published DNaseI hypersensitivity data^54, 55^, our aggregated scATACseq data show that AR significantly altered the chromatin accessibility (**Figure 5A**). This led to an increased regional deviation between each ARBS enhancer class within DHT treated cells (**Figure 5B**). When comparing the relative fold change in accessibility, the largest increase was at inducible enhancers though both inactive and constitutive CRE were significantly altered (**Figure 5C**). As expected, AR binding led to a significant increase in the co-accessibility between ARBS, demonstrating that the AR binds to multiple proximal ARBS following activation (**Figure 5D**). When we incorporated these results into gene-specific networks we observed that gained inducible enhancers frequently increased the number and complexity of interactions between ARBS. A representation of this alteration in co-accessibility is shown in **Figure 5E**. To quantify these changes we used our *cis*-regulatory networks to determine the relative impact of each AR CRE class on network complexity. In agreement with our proposed model, we found that inducible enhancers had the most significant impact on AR CRE interaction complexity and increased interactions with other ARBS (*p<4×10^-4^;* **Figure 5F**). Overall, inducible enhancers significantly increase network interactions between inactive and constitutive CRE and potentially act as a regulatory hub between promoters and other ARBS.

**Figure 5:**
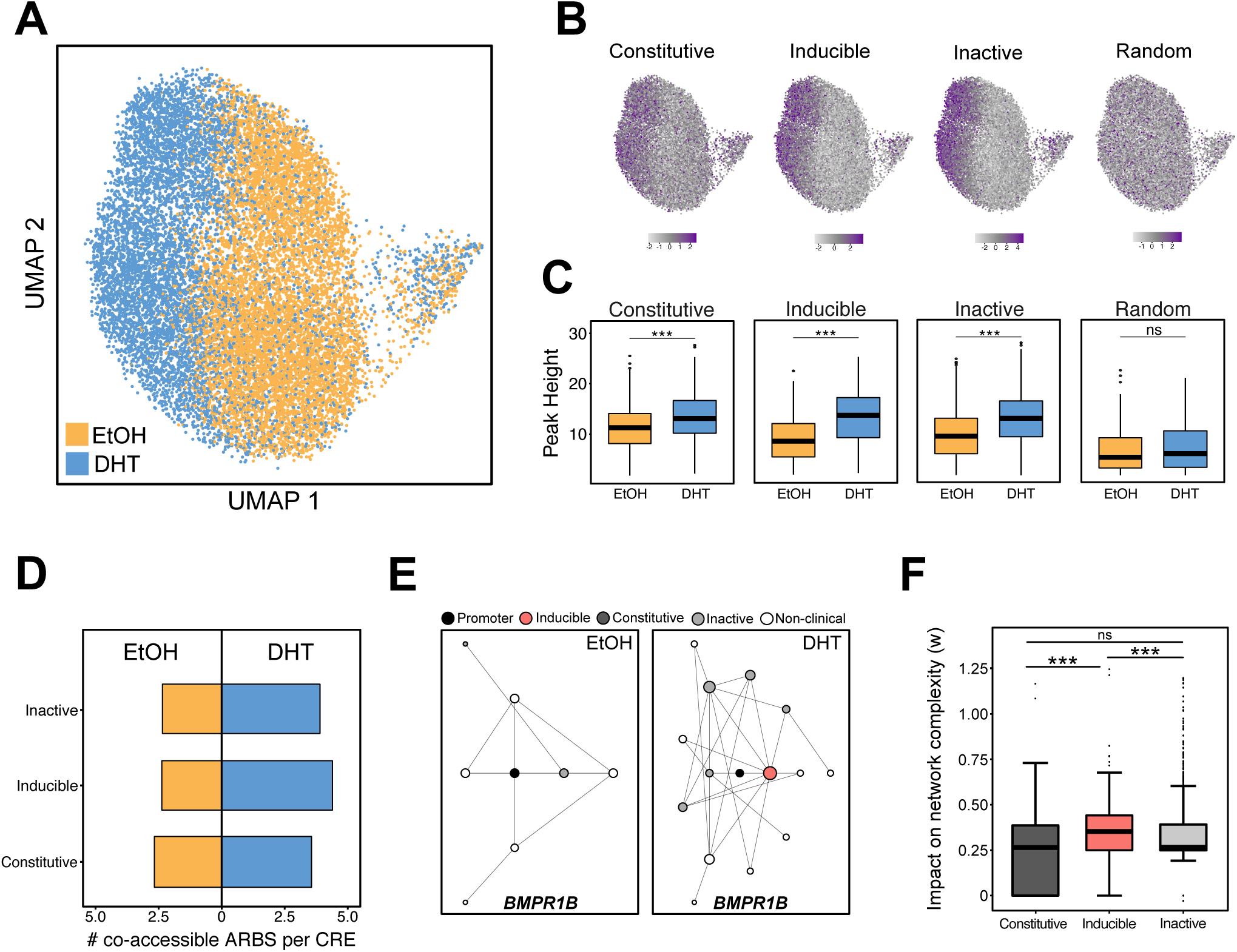
Inducible enhancers are transcriptional ‘hubs’: **(A)** UMAP projection of scATAC-seq profiles of LNCaP cells treated with either ETOH or DHT. Each dot represents an individual cell (EtOH n=7,857; DHT=6,661) **(B)** chromVAR deviation score enrichment of different AR enhancer classifications was compared to random genomic regions in the UMAP projection. (**C)** The chromatin accessibility of pseudo-bulk scATACseq at each AR enhancer class compared to random genomic regions. **(D)** Change in median number of co-accessible sites for each AR enhancer class following androgen treatment. **(E)** A representative network graph (*BMPR1B)* shows changes in co-accessibility following androgen treatment. The gained inducible AR enhancer led to significant increases in complexity. **(F)** The relative impact on network complexity following DHT treatment was calculated for each AR enhancer class. Inducible enhancers had the most significant impact on network complexity leading to higher co-accessibility with other ARBS (*p<10^-6^)*.

### Characterization of genetic alterations

Given the critical function of inducible enhancers on AR-mediated gene expression we speculated that non-coding mutations at these CRE could potentially alter gene transcription in PCa. To identify these potentially important mutations, we overlaid whole genome sequencing of primary PCa (n=196)^56^ and metastatic CRPC (n=101)^20^ with our functional enhancer annotations. As previously published, we observed a significant enrichment of SNVs at ARBS in both primary PCa^23, 24^ and also metastatic CRPC **(Figure 6A)**. Within the annotated ARBS sites we found 751 SNVs in primary PCa and 1013 SNVs in metastatic PCa with 14% of the regions containing overlapping mutations (**Supplementary Figure 13A**). Similar to most protein coding driver mutations in PCa, there were very few recurrent somatic mutations at ARBS^57^. We did not observe any difference in the SNV distribution between inducible, inactive or constitutively active ARBS (Two-sample Kolmogorov-Smirnov test, *p>0.5*). To test the impact of these somatic mutations on AR enhancer activity, we focused on those SNVs in inducible enhancers that looped to the TSS of AR upregulated genes and found that 19% (3/16) of SNVs tested significantly altered AR enhancer activity (**Figure 6B, Supplementary Figure 13B**). Interestingly, one of the affected enhancers was found to interact with the TSS of *ZBTB16,* a well known AR-regulated tumour suppressor that is commonly mutated in CRPC^58, 59^(**Figure 6C**). Critically when this specific AR enhancer was inactivated with CRISPRi we observed a significant decrease in the expression of *ZBTB16* (**Figure 6D**). Taken together, these results show that non-coding SNVs at ARBS can impact enhancer activity of regulatory regions required for the AR-mediated gene expression.

**Figure 6:**
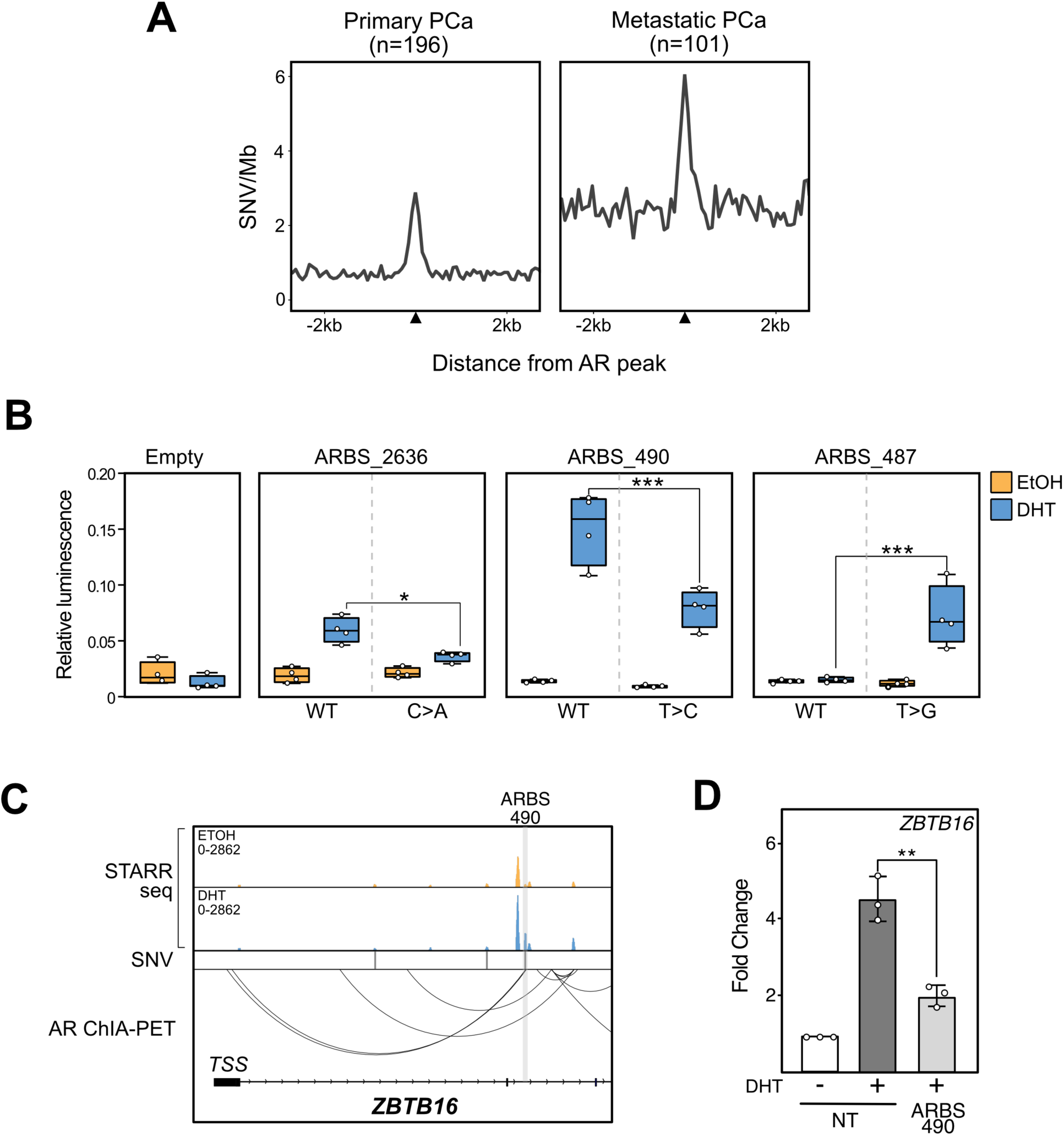
SNVs impact the AR enhancer landscape: **(A)** An increase in SNVs at ARBS was observed in both primary (left) and metastatic (right) PCa. **(B)** The impact of clinical SNV on androgen-dependent enhancer activity was quantified with a luciferase assay (4 biological replicas±SEM; * *p<0.05,* *** *p<0.001*). **(C)** Genome browser snapshot of *ZBTB16* gene locus. Gene looping was observed between enhancer ARBS_490 and *ZBTB16* promoter. **(D)** CRISPRi inhibition of ARBS_490 suppresses androgen-induced expression of ZBTB16 (3 biological replicas±SEM; ** *p<0.01*).

## Discussion

The AR, like most nuclear receptors, binds to thousands of chromosomal sites but only regulates hundreds of genes^49, 60, 61^. The majority of AR-regulated gene promoters therefore physically interact with multiple ARBS^7^. Yet, how these binding sites work together to induce transcription is poorly understood, as annotation of non-coding CRE has been challenging. While descriptive approaches including histone ChIPseq or GROseq largely correlate with enhancer activity, they cannot provide the locus-specific resolution that is needed to understand these complex interactions. To annotate these individual regions we systematically tested the enhancer activity at all commonly occurring clinical ARBS using a MPRA recently optimized for human models^19^. From this we found that only a fraction of ARBS (7%) showed androgen-dependent enhancer activity, while the majority (81%) were inactive. This is analogous to work in stem cells where only a small percentage of CREs marked with NANOG, OCT4, H3K27ac, and H3K4me1 were found to be active enhancers^62, 63^. While the ARBS enhancer usage will likely change during development, the results from this plasmid-based assay strongly correlate with epigenetic features in both PCa cell lines and clinical tumours suggesting that the enhancer capability of the individual CRE are predictive of activity *in situ* (**Figure 1E+2B**). In matched patients pre- and post-ENZA treatment, we observed that H3K27ac at inactive and constitutive enhancers was generally not affected by AR inhibition, while inducible enhancers had a marked decrease when the patient was treated with ENZA. Supporting these results, in a larger patient population of primary PCa samples, H3K27ac was significantly enriched at both induced and constitutively active enhancers as compared to inactive CRE (**Figure 2A**). However, these descriptive ChIPseq results only broadly correlate with AR enhancer activity (**Figure 1G**). They cannot annotate individual CREs and provide the locus-specific resolution needed to characterize complex protein interactions, identify non-coding mutations or delineate the underlying regulatory logic of AR-mediated transcription. By systematically testing all clinical ARBS for enhancer activity, this provides the first detailed “map” needed to investigate these clinically important problems.

Using this functional annotation we then characterized what features correlate with active enhancers. We initially characterized the different DNA motifs in each AR enhancer class but this provided almost no predictive power as ARE motifs were equally distributed between induced, inactive and constitutive enhancers (**Supplementary Figure 9)**. This is in contrast to work with glucocorticoid receptors that suggested steroid response elements were more likely to be found at active enhancers^49^. Potentially this may be due to the larger size of the ARBS tested in our assay, as many motifs will be frequently found in the large fragments (>500bp) incorporated into our STARRseq library. Yet while DNA motifs do not stratify enhancer CREs, AR binding was essential for androgen-mediated enhancer activity. Only those clinical ARBS that had an AR peak in LNCaP cells were inducible enhancers (n=867). Based on these results, we expanded our analysis and trained a machine learning classifier to predict enhancer classes from all publically available ChIPseq studies in LNCaP including transcription factors and histone marks. Once developed, we experimentally validated this model and could successfully identify those regions likely to be inducible AR enhancers (**Figure 3B**). Importantly, as our classifier was not a black box model we could identify those features that were strongly predictive of AR enhancer activity (**Figure 3A**). As this classifier is predictive for AR induction it will not identify those constitutive features found at most ARBS. With this we found that inducible enhancers strongly associate with AR, enhancer-associated features including H3K27ac and MED1, and co-activators such as PIAS1 and ARID1A^6, 64^. When each feature of this larger model was systematically down-sample we found that AR peak height was more predictive of identifying inducible enhancers at ARBS than any feature including H3K27ac (**Figure 3C**). These results help to identify enhancers in those samples that cannot be tested with MPRA but are commonly used for ChIPseq including patient-derived xenografts or clinical samples. While further validation is required, our model supports stratifying AR ChIPseq peaks by height and distinctive co regulatory factors and to identify inducible AR enhancers in samples where functional testing is not possible.

Having annotated the different AR enhancers we then characterized how each class impacted gene transcription. We found that inducible enhancers were more likely than inactive or constitutive enhancers to interact with either other ARBS or TSS of androgen-upregulated genes (**Figure 4A,B+E**). Further, inducible enhancers had the highest number of chromosomal contacts of all ARBS and were central in AR CRE interaction networks (**FIgure 4C+F**). To test their role in transcription we functionally inactivated inducible AR enhancers and found they were critical to androgen-induced transcription (**Figure 4G**). Surprisingly, transcription was not solely dependent on inducible enhancers and that inhibition of constitutive and inactive ARBS frequently impaired androgen-induced transcription. This is unlikely to be a false positive, as we also found that all enhancer classes had similar evolutionary conservation (**Figure 4F**). It is unlikely these CRE would be conserved if they were shadow peaks^65^. While these regions may potentially be inducible enhancers in a different cellular context, our results suggest that these ARBS are involved in gene transcription with multiple classes working in concert to drive transcription. This is similar to estrogen receptor where CRISPRi-mediated knockout of individual binding sites demonstrated that these work in hierarchical or synergistic interactions to induce enhancer-activated gene expression^66^. Further, studies of glucocorticoid receptor-dependent enhancer activity suggested that activation occurs via synergistic interactions to drive the gene expression^49^. Intriguingly, this suggests that enhancer activity is not the only feature required for AR-mediated transcription. Supporting this, AR peak height, a predictor of inducible enhancers (**Figure 3C),** does not strongly predict AR-mediated gene transcription^67^. While speculative, these inactive CRE may work in concert with induced enhancers to increase the local protein concentration and potentially drive the formation of biological phase condensates that have been observed with other nuclear receptors at strongly active enhancers^68^. Supporting this model, we observed that AR binding at inducible enhancers significantly increased co-accessibility between ARBS (**Figure 5F**) which leads inducible AR enhancers to be in close physical proximity multiple ARBS (**Figure 4C**). How these interactions occur is poorly understood and there is conflicting evidence about TF binding inducing new chromatin loops or stabilizing interactions^53, 69^. Regardless of the mechanism, it is clear that inducible enhancers work with other AR CRE to drive AR-mediated gene transcription.

We and others have previously demonstrated that ARBS are highly mutated in primary PCa^23, 24^. Given the critical role of AR in PCa, we speculated that these somatic mutations could potentially alter the transcriptional activity of the tumour and potentially impact PCa growth and proliferation. However, selecting non-coding mutations for experimental validation is challenging due to both the large number of ARBS/mutations and the relatively low frequency of recurrent mutations at CRE^70^. Further compounding this problem, mutations at different CRE can impact gene expression thereby inducing the same phenotype^71^. Yet, while challenging to identify, there is increasing evidence that non-coding somatic mutations play a critical role in PCa disease progression^20–22^. By using our functional enhancer annotation to “map” specific CRE that are likely to impact androgen-mediated transcription we overcome many of these challenges and can select potential somatic mutations for testing. Focusing on induced AR enhancers, we demonstrated that 19% of the SNVs tested significantly altered enhancer activity (3/16; **Figure 6B**). This is significantly higher than a recent study using an orthogonal MPRA of H3K27ac sites which showed 1.8% of primary PCa SNVs impacting enhancer activity^24^. This difference could be due to technical issues related to insert size (146 vs. >500bp), the specific genomic regions tested (H3K27ac vs. ARBS) and the relatively small number of genomic regions tested (n=16). While not comprehensive, of the regions tested approximately 20% of ARBS SNVs significantly impacted enhancer activity. Of particular interest we identified a somatic mutation in metastatic PCa that significantly reduced the activity of a critical AR enhancer required for the expression of *ZBTB16* (**Figure 6D**). This androgen regulated gene is a well characterized tumour suppressor that is frequently mutated in late-stage PCa^58, 72^. Immunohistochemical staining showed a significant reduction of *ZBTB16* in high grade localized PCa specimens and weak or no expression in metastatic PCa biopsies^73^. *ZBTB16* knock-down experiments demonstratedincreased PCa growth^58^ and ectopic expression inhibited prostate cancer tumourigenesis in mouse models^74^. Further, loss of *ZBTB16* promotes a metastatic and ENZA-resistant phenotype in prostate cancer cells^58^. Given such a repressive function, homozygous *ZBTB16* mutations have been found to occur in 4-9% of CPRC tumours^59, 75, 76^. While these mutations have been found to occur in protein coding regions our study demonstrates that somatic mutations in AR enhancer can also reduce *ZBTB16* expression.

In conclusion we have created the first functional map of potential AR enhancer activity. By quantifying the activity of each ARBS, this provides mechanistic insight into mammalian gene regulation. Together our data demonstrates that AR-driven inducible enhancers act as a regulatory hub that frequently cooperate with other ARBS to drive transcription. Identification of key enhancers, stratifies non-coding mutations for functional testing.

## Material and Methods

### Cell lines

Cell lines were purchased from ATCC and routinely tested for mycoplasma contamination. LNCaP cells were routinely grown in RPMI 1640 media (Gibco) with 1% Penicillin-Streptomycin and 10% fetal bovine serum (FBS). No activation of the IFNγ pathway by double stranded DNA was observed in electroporated LNCaP cells which have been shown to systematically alter STARRseq activity^25^. (**Supplementary Figure 1A**).

### Generation of ARBS STARRseq library

Common clinical ARBS were defined as those sites that were present in all normal prostate (n=3) or independent PCa tumours (n=13)^2^. Pooled human male genomic DNA (Promega) was randomly sheared (500-800bp) using a Covaris M220 Focused-ultrasonicator. The fragments were end-repaired and ligated with Illumina compatible adaptors using the NEBNext Ultra^TM^ II DNA Library Prep Kit (NEB). The adaptor-ligated DNA was hybridized to custom Agilent biotinylated oligonucleotide probes across a 700bp region (53032 probes; 4.684 Mbp oligo) and then pulled-down by Dynabeads M-270 Streptavidin beads (NEB). The post-capture DNA library was amplified with STARR_in-fusion_F and STARR_in-fusion_R primers (**Supplementary Table 1**), and then cloned into AgeI-HF (NEB) and SalI-HF (NEB) digested hSTARR-ORI plasmid (Addgene plasmid #99296) with NEBuilder HiFi DNA Assembly Master Mix (NEB). The ARBS STARRseq library was transformed into MegaX DH10B T1R electrocompetent cells (Invitrogen). Plasmid DNA was extracted using the Qiagen Plasmid Maxi Kit.

### ARBS STARRseq

LNCaP cells (>1.3 x 10^8^ cells/replica; 3 biological replicas) were electroporated with 266-300ug (1 million cells:2ug DNA) of the ARBS STARRseq capture library using the Neon Transfection System (Invitrogen). Electroporated cells were immediately recovered in RPMI 1640 medium supplemented with 10% FBS. After overnight recovery, the media was changed to RPMI 1640 medium supplemented with 5% charcoal stripped serum (CSS), and 1% Penicillin-Streptomycin. Approximately 72 hours after electroporation, the cells were treated with 10nM DHT or EtOH for 4 hours, washed with PBS and then lyzed using the Precellys CKMix Tissue Homogenizing Kit and the Precellys 24 Tissue/Cell Ruptor (Bertin Technologies). Total RNA was extracted using Qiagen RNeasy Maxi Kit (Qiagen) and the mRNA was isolated using the Oligo (dT)25 Dynabeads (Thermo Fisher). Isolated mRNA samples were treated with Turbo DNase I (Thermo Fisher), synthesized into cDNA using the the gene-specific primer (**Supplementary Table 1**), treated with RNaseA (Thermo Fisher), and PCR-amplified (15 cycles) with the junction PCR primers (RNA_jPCR_f and jPCR_r primers, **Supplementary Table 1**). The ARBS STARRseq capture library was PCR amplified with DNA specific junction PCR primers (DNA_jPCR_f and jPCR_r primers, **Supplementary Table 1**). After junction PCR and AMPure XP beads clean-up, an Illumina compatible library was generated by PCR amplification with TruSeq dual indexing primers (Illumina) and sequenced on Illumina HiSeq4000 (150bp; PE). The resulting sequencing data is available at GSE151064.

### Analysis of STARRseq data

Reads were mapped to reference genome (hg19) with BWA aligner (v0.7.17)^26^ and all mapped reads with a MAPQ score <60 or indels were removed. Captured region coverage was quantified with BamCoverage function (v3.1.3) in DeepTools^27^ while discarding all reads on blacklisted regions (ENCODE ENCFF001TDO). Differential enhancer activity was quantified by DESeq2 (v1.26.0)^28^. Induced enhancers were defined as having a log2 fold-change (LFC)>1 and *p*-adj<0.05 in the DHT/EtOH samples. Constitutively active enhancers had plasmid-normalized reads LFC>1 in both EtOH and DHT but no DHT induction (DHT/ETOH LFC<1). Inactive regions had both minimal DHT inducible activity and low plasmid to RNA ratios (plasmid-normalized LFC<1). Output of the DESeq2 was visualised with ggplot2^29^.

### Clinical approval and sample collection

Clinical PCa tissue was collected before and after Enzalutamide (ENZA) therapy from the Dynamics of androgen receptor genomics and transcriptomics after neoadjuvant androgen ablation study (ClinicalTrials.gov #NCT03297385). The trial was approved by the IRB of the Netherlands Cancer Institute. Informed consent was signed by all participants enrolled in the study and all research was carried out in accordance with relevant guidelines and regulations. Trial participants received three months of neoadjuvant ENZA treatment prior to robotic-assisted laparoscopic prostatectomy. Biopsy (pre-treatment) and prostatectomy specimens (post-treatment) were fresh-frozen and sectioned prior to immunoprecipitation. Tissue sections were examined pathologically for tumour cell content and only those samples with a tumour cell percentage of ≥50% were used for tissue ChIPseq analysis.

### Tissue ChIPseq

ChIP on prostate cancer biopsy- and prostatectomy-tissues were performed as previously described^30^. Nuclear lysates of each tissue specimen were incubated with 5 μg of H3K27ac antibody (Active Motif, 39133) pre-bound to 50 µL magnetic protein A dynabeads (Thermo Fisher Scientific, 10008D). Immunoprecipitated DNA was processed for library preparation (Part# 0801-0303, KAPA biosystems kit) and samples were sequenced using an Illumina HiSeq 2500 system (65 bp, single-end). Sequences were aligned to the human reference genome hg19, duplicate reads were removed, and reads were filtered based on MAPQ quality (≥ 20).

### Tissue ChIPseq data processing

Intensity plots were generated using EaSeq^31^. For boxplots, the number of sequence reads per region of interest was calculated using bedtools multicov (v2.25.0)^32^. The data was further processed in R (v3.4.4) (R Core Team, 2018). Region read counts were z-transformed per sample to correct for differences in total read count. Statistical significance in read counts differences was determined using the Mann-Whitney test, based on the median read count over all samples, and adjusted for multiple testing using FDR.

### RNAseq and ChIA-PET analysis

AR-regulated genes were identified from publicly available RNAseq data of LNCaP cells treated with 100nM DHT for 6 hours (GSE64529) with DESeq2 (v1.26.0)^28^. The distance between an androgen upregulated DEGs and ARBS was calculated with HOMER (v4.10)^33^. The resulting data was merged in 100bp bins and the cumulative distribution function was determined with *Ecdf* in R (v3.6.1). Specific ARBS-promoter interactions were identified from the published AR ChIA-PET (GSE54946) by overlapping the loop end of either ARBS or gene’s TSS (+/- 5Kb) with GenomicInteractions (v1.20.3)^34^ .

### ChIPseq and GROseq analysis

Previously published ChIPseq and GROseq data were downloaded from the GEO database and uniformly processed (**Supplementary Table 2).** The sequencing reads were controlled for quality using FASTQC and the reads were then mapped to the human reference genome (hg19) with bowtie aligner (v0.12.9)^35^. All reads mapped to the blacklisted regions (ENCODE ENCFF001TDO) were discarded. For direct comparison of the H3K27ac ChIPseq with the STARRseq data, average signal values of the inducible and inactive regions were calculated using bigWigAverageOverBed software(v377)^36^. For each region, log fold change values were calculated between DHT and ETOH treatments in H3K27ac and STARRseq experiments. Scatterplots were generated by R’s ggplot2 package (v3.2.0)^29^. ROSE^37^ was used to identify super enhancers from DHT H3K27ac ChIPseq. BigWig signal tracks were generated with BamCoverage function (v3.1.3) of deepTools^27^ with RPKM normalization. For GROseq data, using log fold enrichment between + strand of DHT over the + strand EtOH a single track was generated. Similarly a single - track was also generated. Then - strand is subtracted from + strand to final bigwig file. For each of the 500 positive control enhancer regions, we calculated the average accessibility scores using bigWigAverageOverBed from ENCODE’s LNCaP DHT (ENCFF975MZT) and EtOH (ENCFF906QXX) DNase-seq experiments. Then we compared the top and bottom 100 to show differences in accessibility. Later, for the same regions we calculated the average STARRseq signal with bigWigAverageOverBed^36^ from merged DHT and EtOH samples.

### Conservation

100 vertebrate species conservation track was obtained from UCSC golden path (http://hgdownload.cse.ucsc.edu/goldenpath/hg19/phastCons100way/hg19.100way.phastCons. bw) for hg19 reference. The average conservation distribution of induced, constitutive, inactive ARBS enhancers and negative control regions were calculated with deeptools’ *computeMatrix* function (v3.1.3). Output distribution matrix then visualised with deeptools’ *plotProfile* function (v3.1.3).

### Network Graph of AR CRE

A network graph was built using the annotated CREs (clinical ARBS and TSS) and LNCaP ARBS (not tested for enhancer activity) as nodes and the chromosomal interactions from VCaP ChiA-PET as edges. The interactions between each enhancer class or ARBS were extracted from processed data and introduced as a graph with NetworkX (v2.3). Only TSS of AR DEG were considered in the network. The annotated CRE network graph contained 2361 nodes, 1951 edges and 830 separate components. The annotated CRE and non-annotated ARBS graph contained 10484 nodes, 12437 edges and 1873 separate components. The relative density (eq.1) of two different classes was calculated based on the expected frequency between two classes. If self-looping occurred within the same class the expected maximum edge frequency was duly calculated to reduce duplication (eq.2). For different classes in the bipartite graph the expected maximum edge frequency is calculated accordingly (eq.3). The interaction frequency (eq.4) was scored as the ratio of observed versus expected edge number relative to whole graph density.

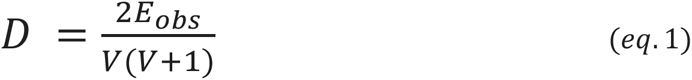

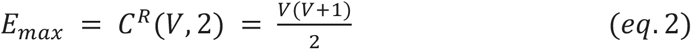

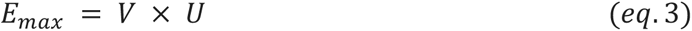

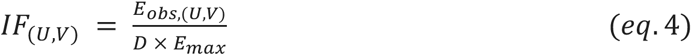

Degree centrality and betweenness centrality scores were calculated by NetworkX. Although a not-connected graph gives information about the whole network, it is biased when comparing the centrality features of all elements. Therefore, we calculated the centrality score from the largest connected component.

### Enhancer luciferase assay

The region of interest (750-850 bp) was PCR amplified from pooled male human DNA (Promega) and cloned into a STARRseq luciferase validation vector_ORI_empty plasmid (Addgene plasmid #99298) with Hi-Fi DNA builder (NEB). Primers used to amplify specific regions are described in (**Supplementary Table 1**). LNCaP cells were co-transfected with 500ng reporter plasmid and 5ng of Renilla using TransIT-2020 (Mirus) and plated in phenol red-free RPMI (Gibco) supplemented with 5% CSS (Fisher Scientific). 48h post-transfection, cells were treated with 10nM of DHT or ETOH for 24 hours. Firefly and Renilla luciferase activity were assayed by Dual-Glo Luciferase assay system (Promega). All the experiments had a minimum of 4 biological replicas with 3 technical replicates in each experiment. Single nucleotide substitutions of reporter constructs were carried out by inverse PCR mutagenesis as previously described^38^. All mutagenic primers are given in (**Supplementary Table 1**).

### gRNA design and CRISPRi

Multiple gRNAs per enhancer region were designed with CRISPR-SURF^39^ and cloned into lentiGUIDE-puro (Addgene #52963). All gRNA sequences are provided in **Supplementary Table 1**. LNCaP cells stably expressing dCas9-KRAB were generated by transducing LNCaP cells with Lenti-dCas9-KRAB (Addgene #89567) followed by blasticidin selection. dCas9-KRAB expression was confirmed by Western Blot (Cas9: CST Mouse mAB #14697, GAPDH: SC Rabbit pAB #25778) (**Supplementary Figure 1B)**. LNCaP-dCas9-KRAB cells (200,000/well) were transfected with 1µg of gRNA using Mirus TransIT-X2 and selected with puromycin (2µg/ml) for 72 hours. The media was then changed to phenol-red free RPMI supplemented with 5% CSS and treated with EtOH or 1nM DHT for 24 hrs. Androgen induced expression was quantified by qRT-PCR using gene specific primers (**Supplementary Table 1**). As CRISPRi is known to be prone to false negatives, all gRNA were initially tested and then a single gRNA was used against each genomic region. In each set, expression of the non-AR regulated gene *FXN* was also quantified to assess non-specific inhibition. Each experiment was done in triplicate with a minimum of 3 biological replicas.

### Single cell ATACseq (scATACseq)

LNCaP cells were cultured in phenol red-free RPMI 1640 media supplemented with 5% CSS and 1% antibiotics for 60 hours and then treated with 100% EtOH or 10nM DHT for 4 hours. Nuclei from 1×10^6^ cells were isolated according to 10x Genomics recommended protocol (Nuclei Isolation for Single Cell ATAC Sequencing CG000169 Rev D). scATACseq libraries were prepared using 10x Genomics Chromium Next GEM Single Cell ATAC Library & Gel Bead Kit v1.1. Libraries were sequenced on a NextSeq500 Illumina sequencer (23,428 unique PE reads/cell DHT and 27,077 unique PE reads/cell EtOH). scATACseq analysis was done with 10X Cell Ranger software. Deviation/z-score was calculated with chromVAR^40^ and UMAP was calculated in ArchR^41^ (v0.9.3). ATACseq peaks were called using MACS (v2.2.6)^42^. Random genomic regions were generated using bedtools (v2.29.2) ^43^. scATACseq co-accessibility scores in EtOH and DHT treated LNCaP scores were calculated using Cicero (v1.4.0)^44^ with 10X Genomics input. Co-accessible sites overlapping with STARRseq annotated AR enhancers and LNCaP ARBS (GEO: GSM2219854) were calculated with bedtools (v2.29.2)^32^ and counted with R (v3.6.1). With these results, two network graphs (EtOH+DHT) were built with annotated AR CREs, LNCaP ARBS (not present in STARRseq library) and promoters as nodes with the predicted Cicero interactions as edges (co-accessibility score >0.1). Inexact graph matching was used to calculate the similarity score of these networks with the DeltaCon, a massive-graph similarity function algorithm^45^. The *S* affinity matrices of the was calculated as described (eq.5) where *I* is the identity matrix, *D* is the diagonal degree matrix, and *A* is the adjacency matrix of *G* and *ε* is the constant defined as 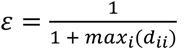. Next the Euclidean distance of two affinity matrices (*d*) was calculated (eq.6).

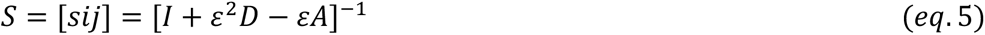

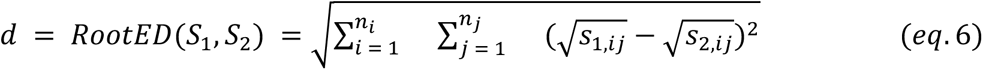

If an edge adjacent to the node (*v*) was altered the impact (*w*) was calculated as the *RootED* distance between corresponding row vector *S*_1,*v*_ and *S*_2,*v*_ calculated where *v* = *j* = 1 → *n_j_* and *n_i_* = 1.Otherwise, *w* = 0.

### Machine learning of ARBS activity

All ChIPseq data used in machine learning was analysed with the standardized ChIP-Atlas bioinformatic pipeline^46^. Based on the inducible (N=286), inactive (N=2479) and constantly active (N=465) categorization of the clinical ARBS we trained a classifier to predict the groups using the bound factors in a given region as input. For each ARBS region, we extracted the ChIPseq signal scores over a 750 bp region for 90 different DNA binding factors (**Supplementary Table 2**). To correct for variations in scores across factors and to unify their values to a consistent range, we applied SES normalization to estimate a score cut-off that separates non-specific from specific binding in each ChIPseq dataset^47^. Briefly, the method finds the score where the difference between the cumulative distributions of the observed and control input scores are maximally different. The median value of the scores above the binding cutoff was then used as the mean of a sigmoid transformation for each bound factor to transform individual factors to an occupancy score between 0 and 1. Heatmaps of the average occupancy score for each bound factor at a 50 bp resolution for inducible and inactive enhancers are shown in **Supplementary Figure 2A+B**. Finally, we took the maximum occupancy score over the 750 bp region as the feature of the factor’s activity. The classifier we chose to fit was a bootstrapped multinomial logistic regression model with a sparsity LASSO regularizer. (Several other regularizations were tried including Ridge and Elastic) but this was found to give the best accuracy and interpretability). In an attempt to balance the number of samples between groups, we created dataset samples that consisted of 500 randomly selected samples of the Non-Inducible group alongside all of the samples from the Constitutively Active and Inducible ARBS groups. The data set was further split into 80 percent training and 20 percent testing. The bootstrapped model consists of 100 thousand different base logistic regression estimators which were all trained on a different subset of the training data. Each subset contains a maximum of 50 training samples and 5 occupancy features. The fitted weights in the aggregate model are an average of the fitted weights in the base estimators and reflect the importance of a factor to differentiate the categories in the presence and absence of other factors. This eliminates the problems of collinearity and the linear dependence within our features and allows for a robust representation of feature importance. Assessing these classifiers showed both false negatives (**Supplementary Fig. 2C**) and positives (**Supplementary Fig. 2D**). False negatives were likely due to missing values in the ChIPseq dataset as this group was much more likely to have a missing value than the inducible group as a whole. False positives were still mostly above zero, indicating potential enhancer function that fell just below the fold-change cutoff to be considered in the inducible group (**Supplementary Figure 2D**). To rank the different factors in terms of predictive power for a given group, we computed the average binding energy of a given factor as the fitted weight for that group times the average occupancy for that factor over all regions in that group. The difference in binding energies between groups could then be used to identify features that differentially predict one group over another. For the 10000 LNCaP specific ARBS regions, we computed their occupancy and constructed their 90 input features as above. The trained classifier was then used to predict the probability of each of the three categories for each of these regions using their occupancy features as input.

## Supporting information

https://docs.google.com/document/d/1w6qPf-SBIF2m9dmDQkOKcDi1sZAj6AMut8emfuFr7hE/edit

## Acknowledgement

We would like to acknowledge the NKI-AVL Core Facility Molecular Pathology & Biobanking (CFMPB) for supplying biobank material and lab support, the NKI Genomics Core Facility for Illumina sequencing and bioinformatics support, and the NKI Research High Performance Computing (RHPC) facility for computational infrastructure. This work was supported by funding from KWF Dutch Cancer Society (10084 ALPE), TUBITAK 1001 (119Z279), NSERC, Prostate Cancer Foundation BC and Astellas Pharma Inc.

---

**Supplementary Figure 1:**
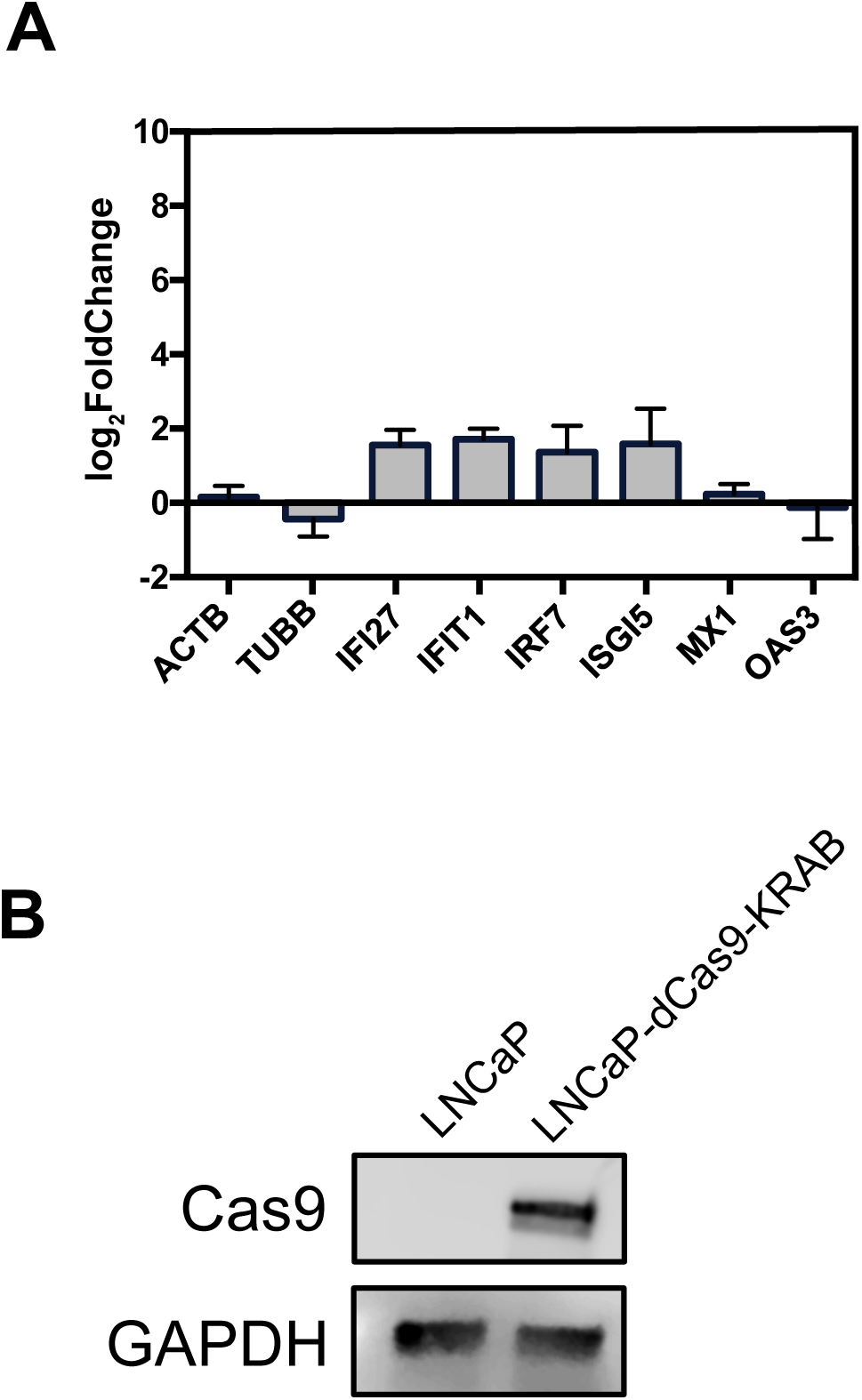
**(A)** The expression of genes involved in the IFNγ signaling pathways quantified by qRT-PCR in LNCaP cells transfected with the ARBS STARRseq library. The bar graph shows the mean expression with standard-deviation from 3 biological replicates. **(B)** Lysates form LNCaP cells with and without stably expressing dCas-KRAB were separated on SDS-PAGE and probed with antibody against Cas9. B-actin was used as the loading control.

**Supplementary Figure 2:**
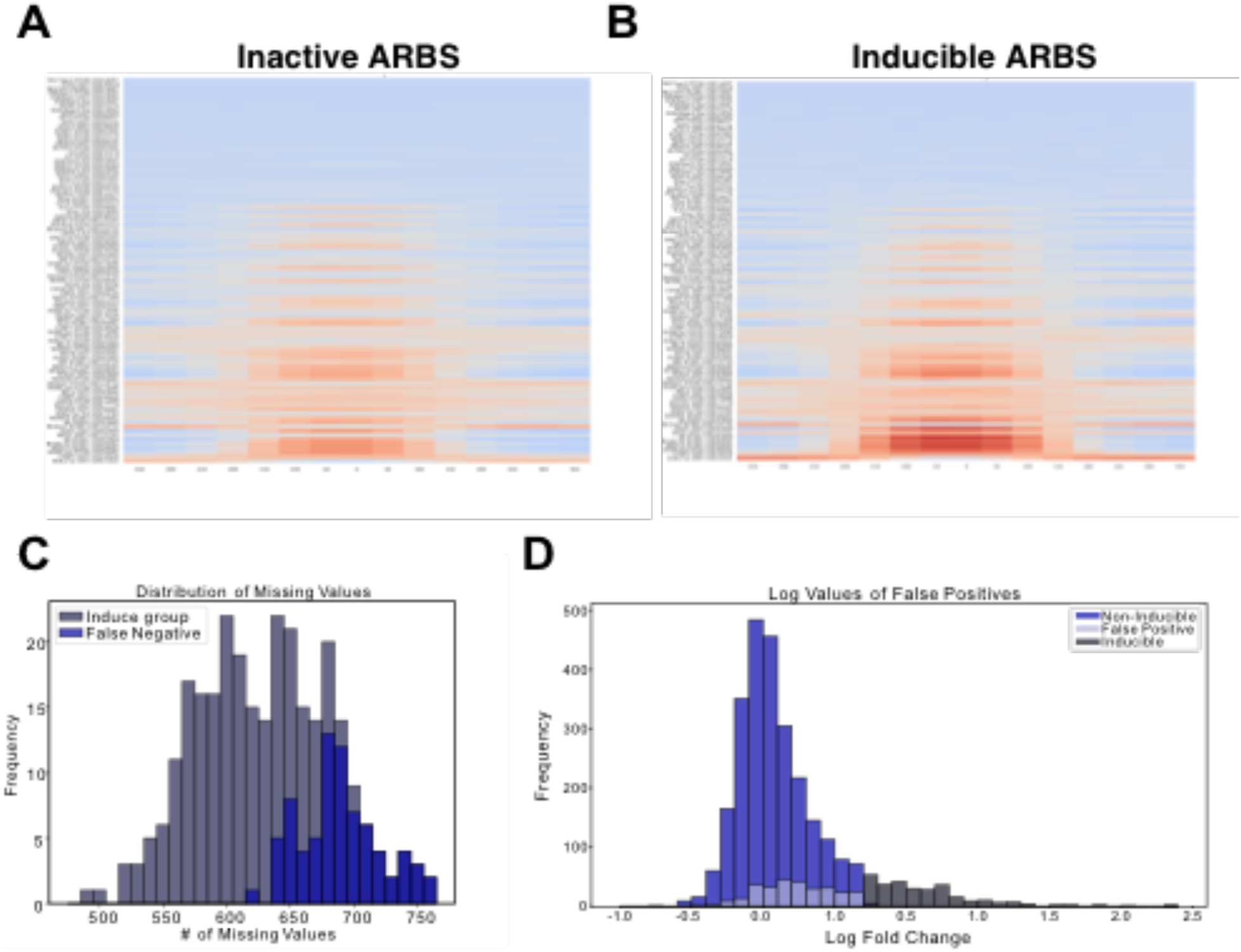
Heatmap of average occupancy score (50 bp resolution) for inducible and inactive AR enhancers in EtOH **(A)** and DHT **(B)** treated LNCaP cells. Factors with a below-average occupancy score are shown in blue while those with an above-average occupancy score are shown in red. **(C)** Distribution of missing values in ChIPseq data for all inducible regions and those misclassified inducible regions. The x-axis represents the number of missing values within a 50bp region. Misclassified inducible regions have a larger proportion of missing values, leading to poorly defined inputs to the classifier. **(D)** Distribution of androgen induced STARRseq expression for regions that were classified as inactive, inducible, and falsely predicted to be inducible by the classifier. The distribution is both skewed towards positive values and close to the cutoff for the inducible group leading to incorrect predictions.

**Supplementary Figure 3:**
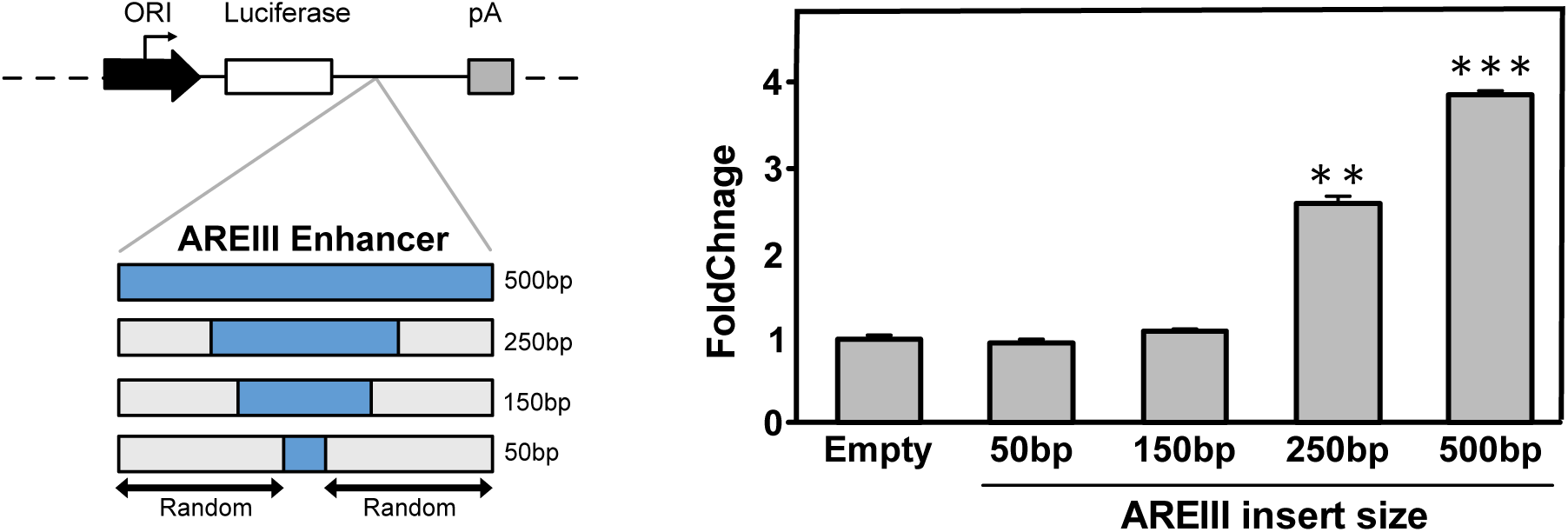
ARBS size is critical for its enhancer functions. A fixed insert size was cloned into the modified STARR ORI luciferase plasmid with varying AREIII enhancer and random sequences (left). Androgen-activated luciferase activity of these size-matched constructs (3 biological replicate±SD; ** *p< 0.01*). (Right)

**Supplementary Figure 4:**
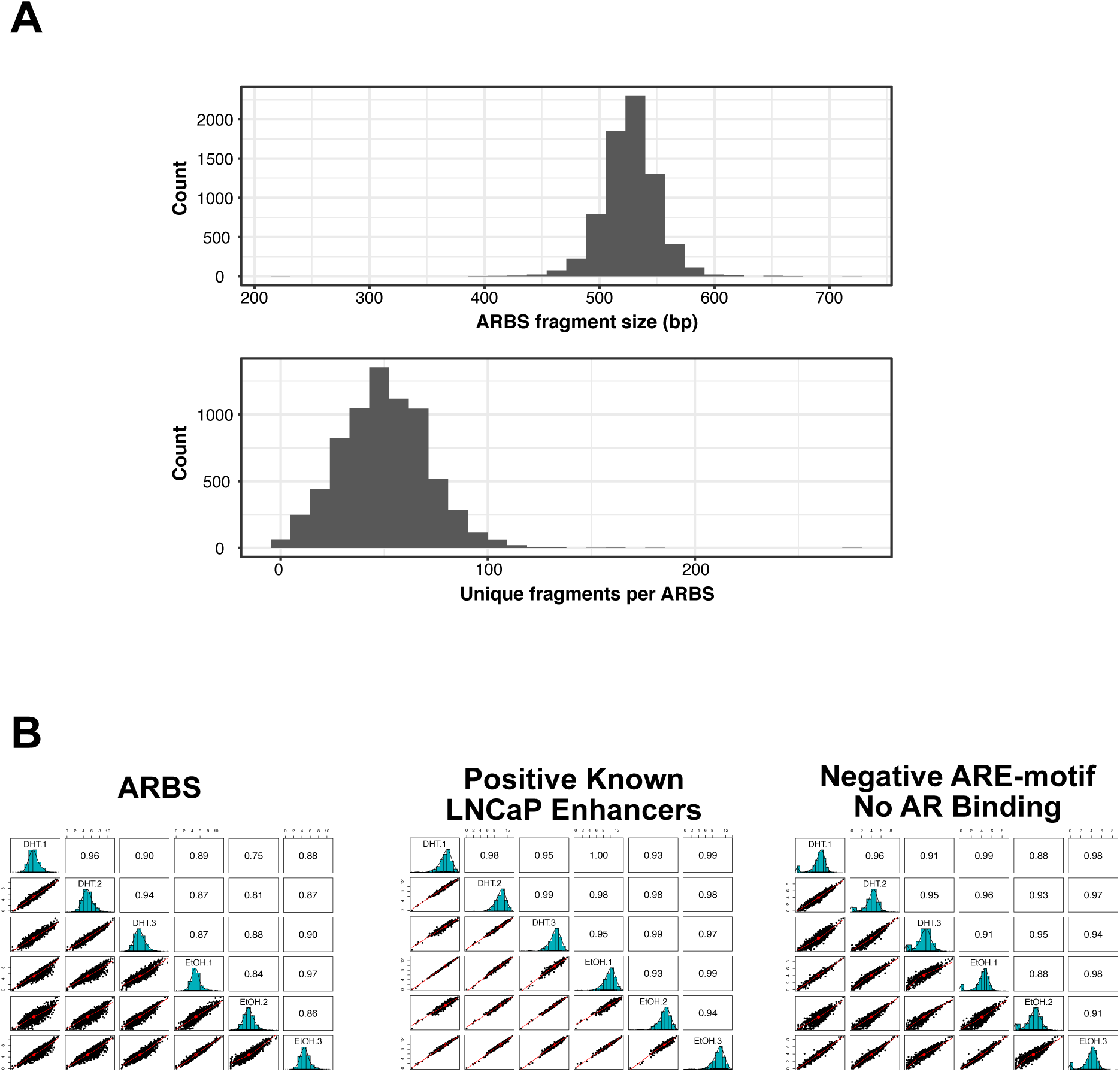
**(A)** Clinical ARBS were captured from normal genomic DNA using a custom-capture and cloned into STARRseq plasmid library. STARR-seq shows a normal distribution of the fragments covering the whole of the capture region. **(B)** Pearson correlation of count data between three biological replicas in each ARBS class.

**Supplementary Figure 5:**
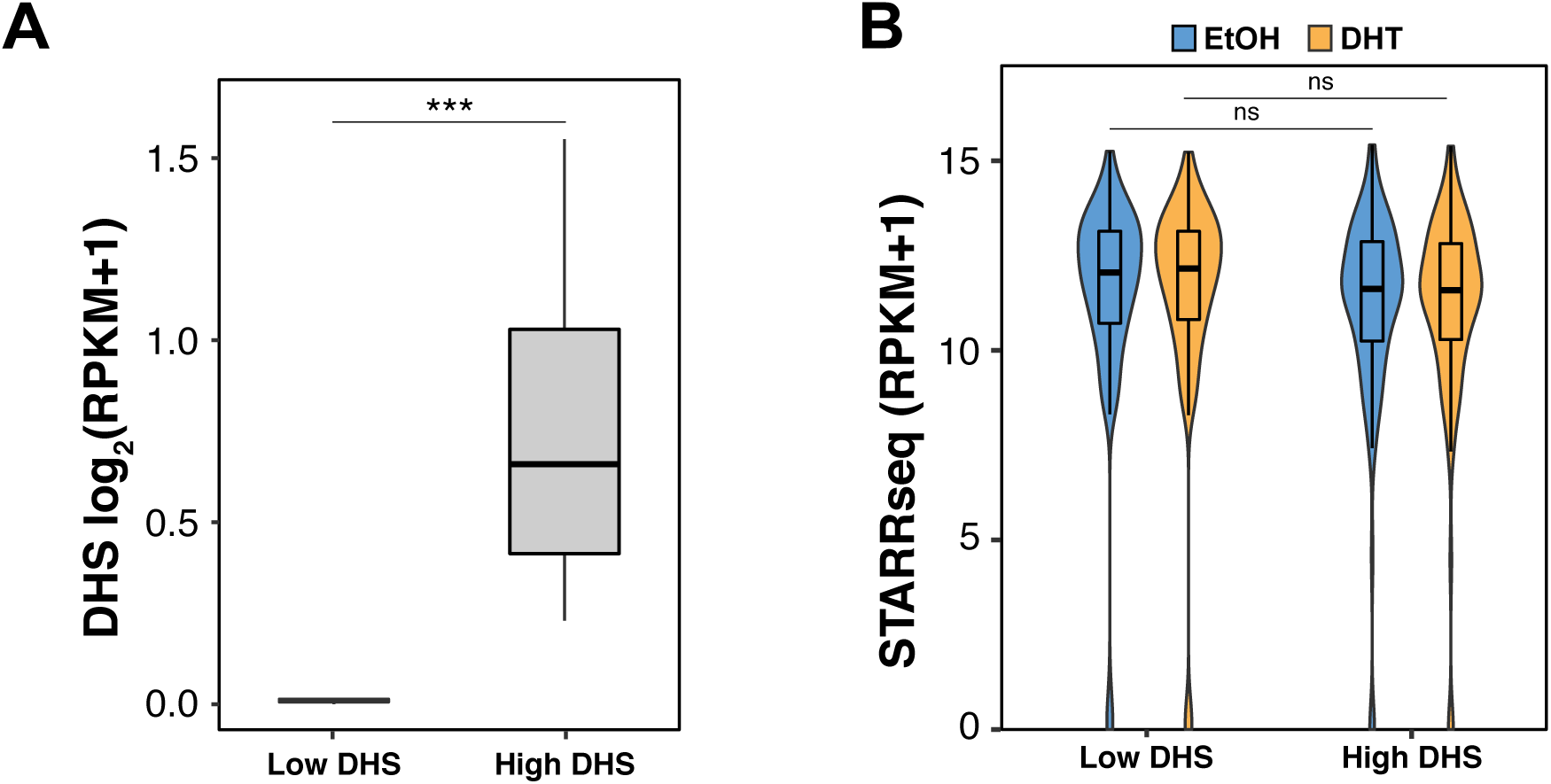
**(A)** Non-AR LNCaP enhancers (positive control) were divided into two groups based on their chromatin compaction in LNCaP cells; heterochromatin (n=100; low DHS) or euchromatin (n=100; high DHS). **(B)** The violin plot shows the enhancer activity quantified by STARR-seq the regions with low DHS and high DHS. Similar enhancer activity is observed in both the groups regardless of the endogenous chromatin compaction.

**Supplementary Figure 6:**
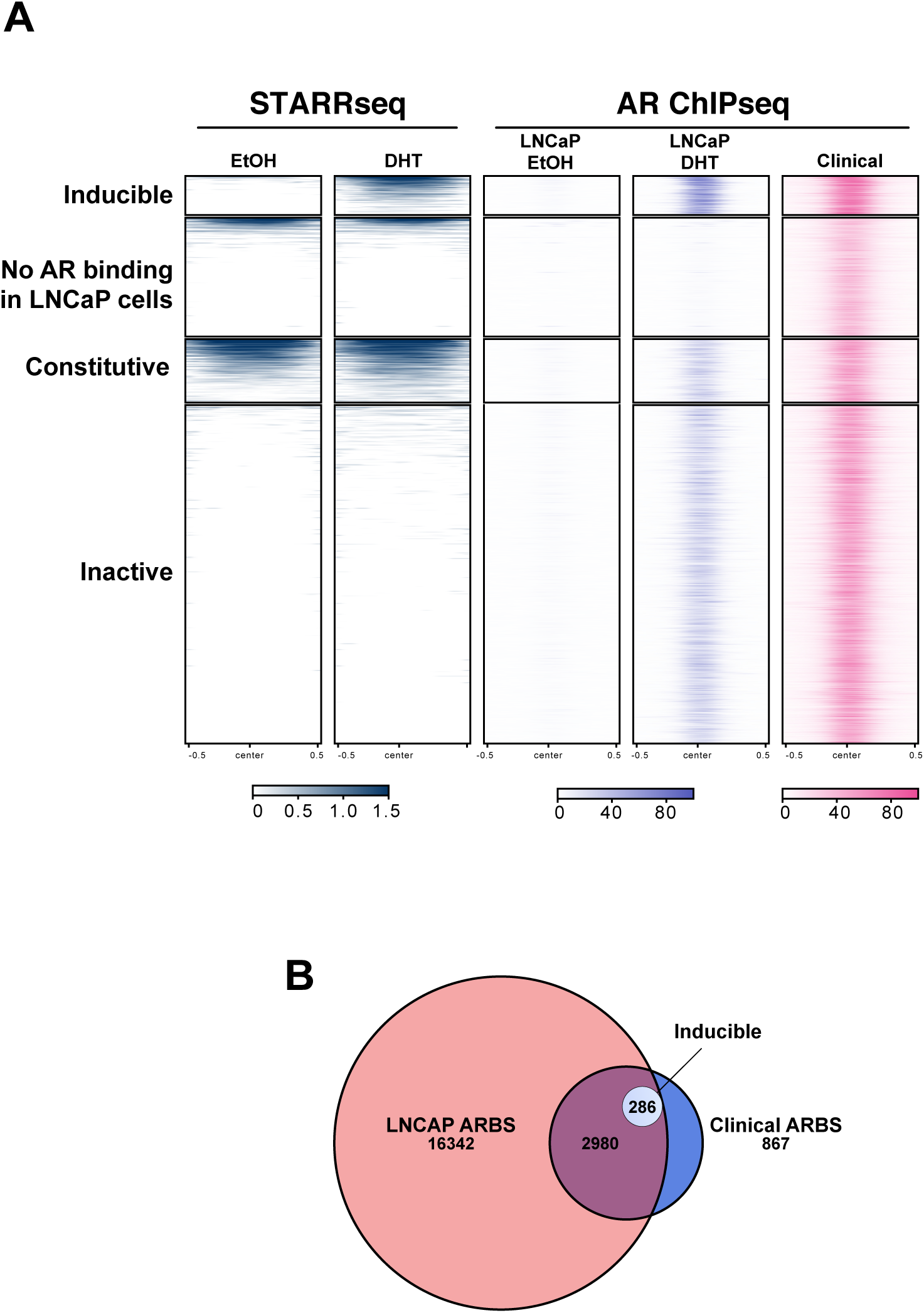
**(A)** Heatmap of STARR-seq and AR ChIPseq in LNCaP cells and primary PCa samples (n=13). None of the ARBS found only in clinical samples show enhancer activity in LNCaP (n=900). **(B)** Venn diagram shows the overlap of ARBS in clinical samples and LNCaP cells. Only those ARBS common to both clinical samples and LNCaP cells show androgen-dependent enhancer activity.

**Supplementary Figure 7:**
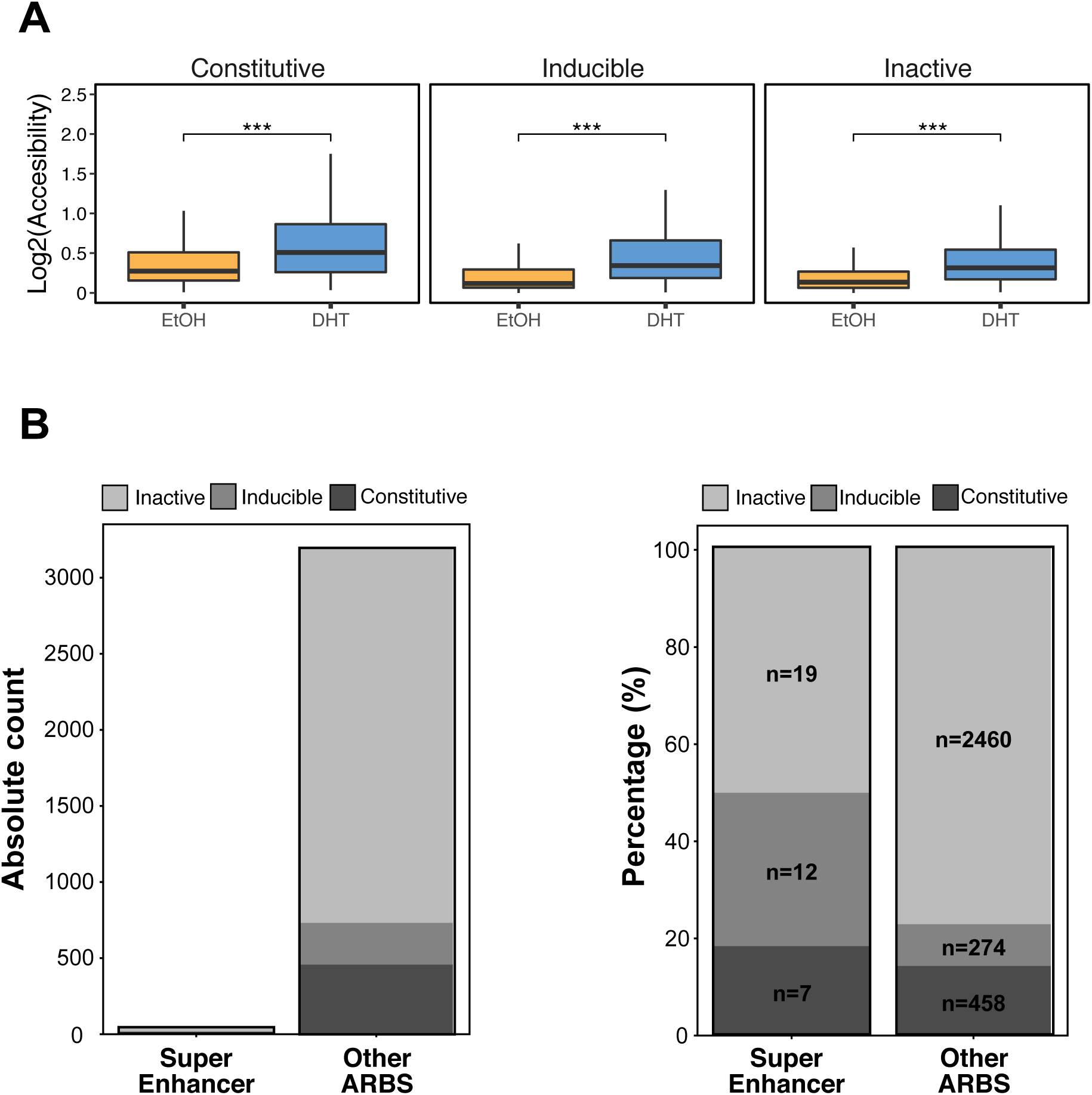
**(A)** DHS of each enhancer class +/- DHT treatment in LNCaP cells **(B)** The total number of super enhancers found in each AR enhancer class of (Left) and their relative percentage (Right).

**Supplementary Figure 8:**
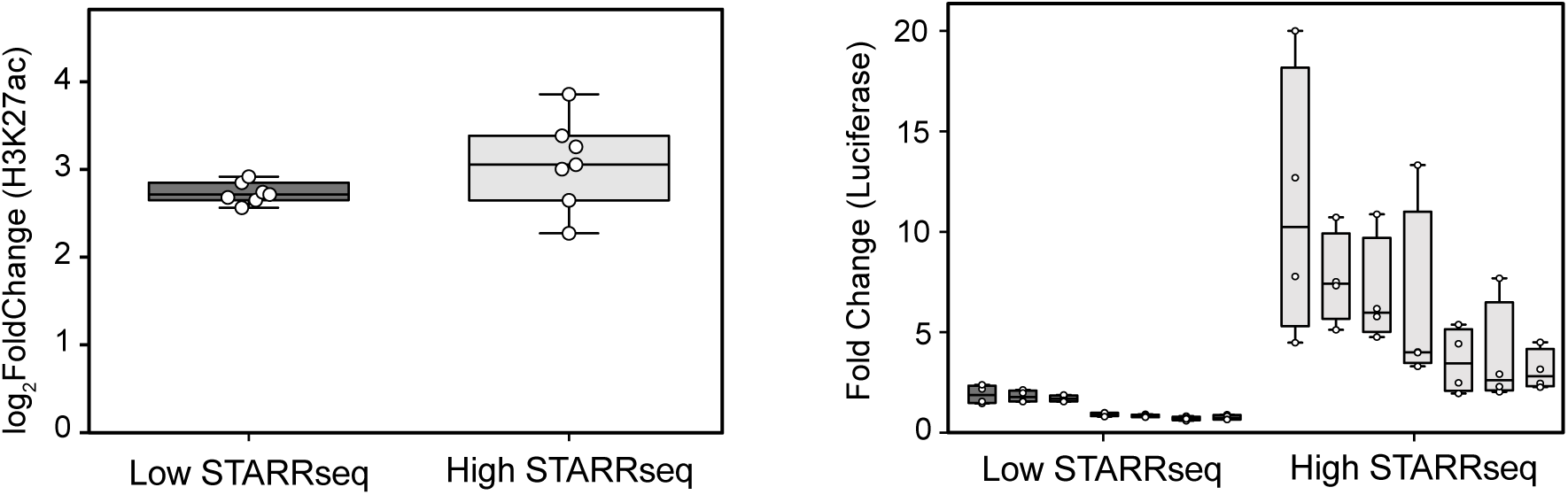
The enhancer activity of ARBS with high H3K27ac and either low inducible (n=7) or high inducible STARRseq signal (n=7) were validated by traditional luciferase assay. While both the groups showed high H3K27ac (left) only the high inducible STARRseq group had androgen-induced reporter activity with a luciferase assay (right).

**Supplementary Figure 9:**
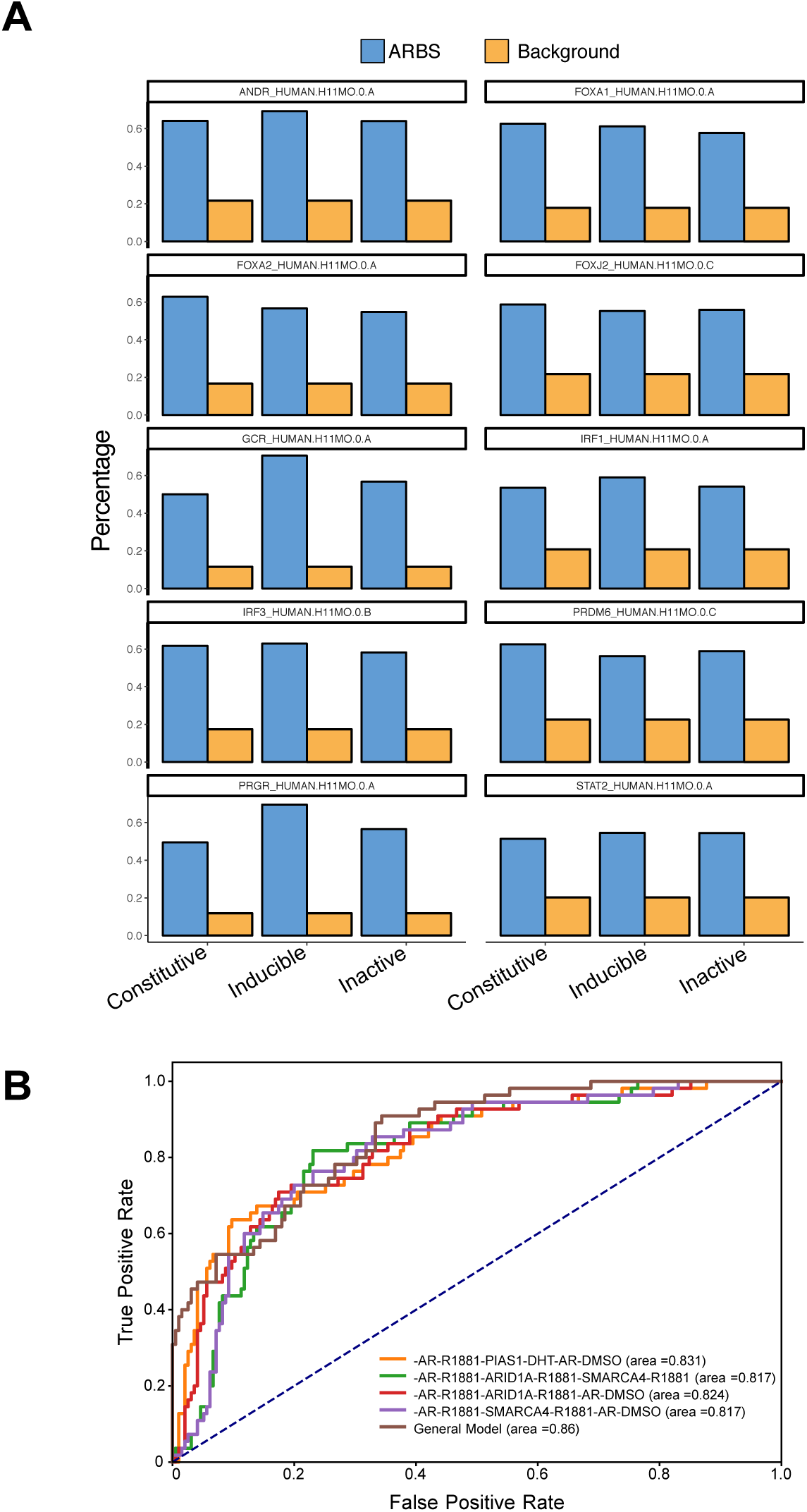
**(A)** Motif analysis was performed to identify the features associated with each enhancer class. The graph shows the percentage of enrichment of indicated motif in relation to the background in all 3 different classes of enhancers across the most highly enriched motifs (n=10). **(B)** Receiver operating characteristic curve of the top three downsampled features (4/117480 permutations) compared to the larger general model.

**Supplementary Figure 10:**
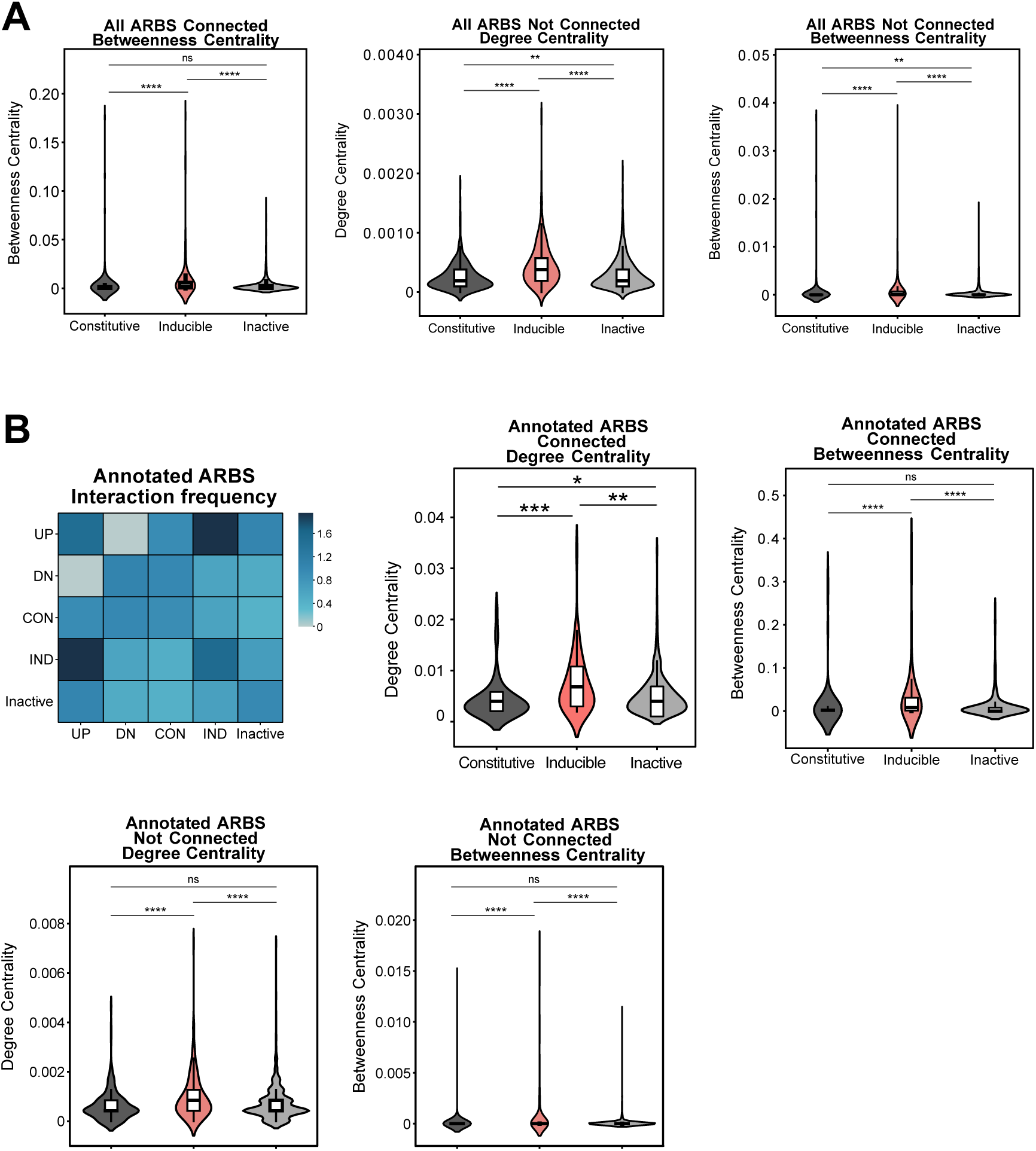
Interaction network of clinical ARBSs. **(A)** In the largest connected network the betweenness centrality score is shown for all AR enhancer classifications. **(B)** For the complete network the degree centrality score (left), and betweenness centrality scores (right) are shown for all AR enhancer classification. **(C)** In an interaction network of only annotated clinical ARBS the interaction frequency between each node class (top-left), degree centrality scores in the biggest connected component (top-middle), betweenness centrality scores in the biggest connected component (top-right), degree centrality scores for the whole network (bottom-left), and betweenness centrality score for the whole network (bottom-right). (ns *p>0.05, **** p<10^-9^*)

**Supplementary Figure 11:**
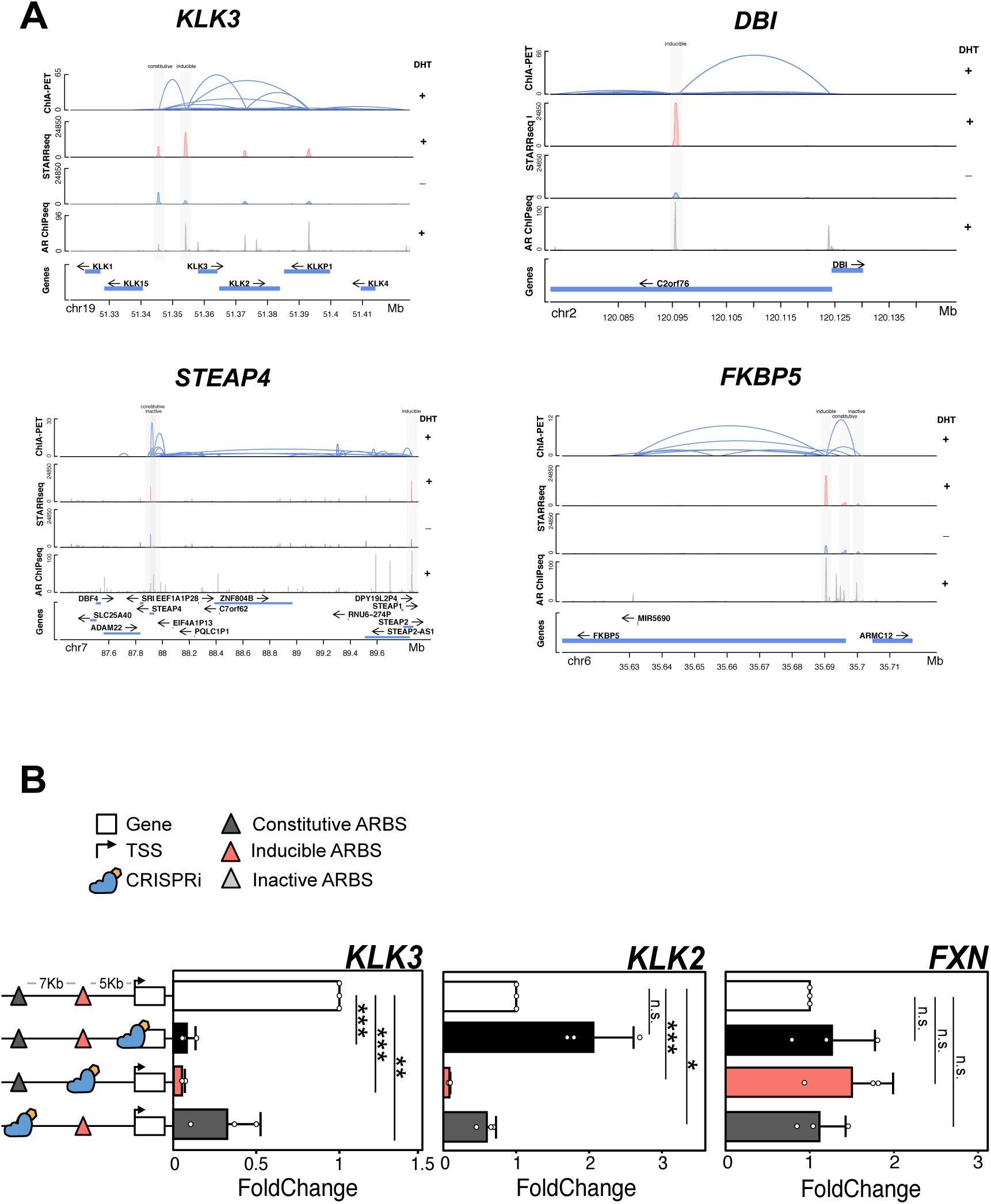
**(A)** Genome browser snapshot of cis-regulatory regions around characterized AR-regulated genes. Long-range chromatin interactions are shown in blue. STARR-seq peaks are shown in red and AR ChIP-seq peaks in grey. **(B)** DHT mediated activation of KLK3, KLK2 and FXN were quantified by qPCR following CRISPRi targeting either KLK3 promoter or KLK3 enhancers (3 biological replica±SD).

**Supplementary Figure 12:**
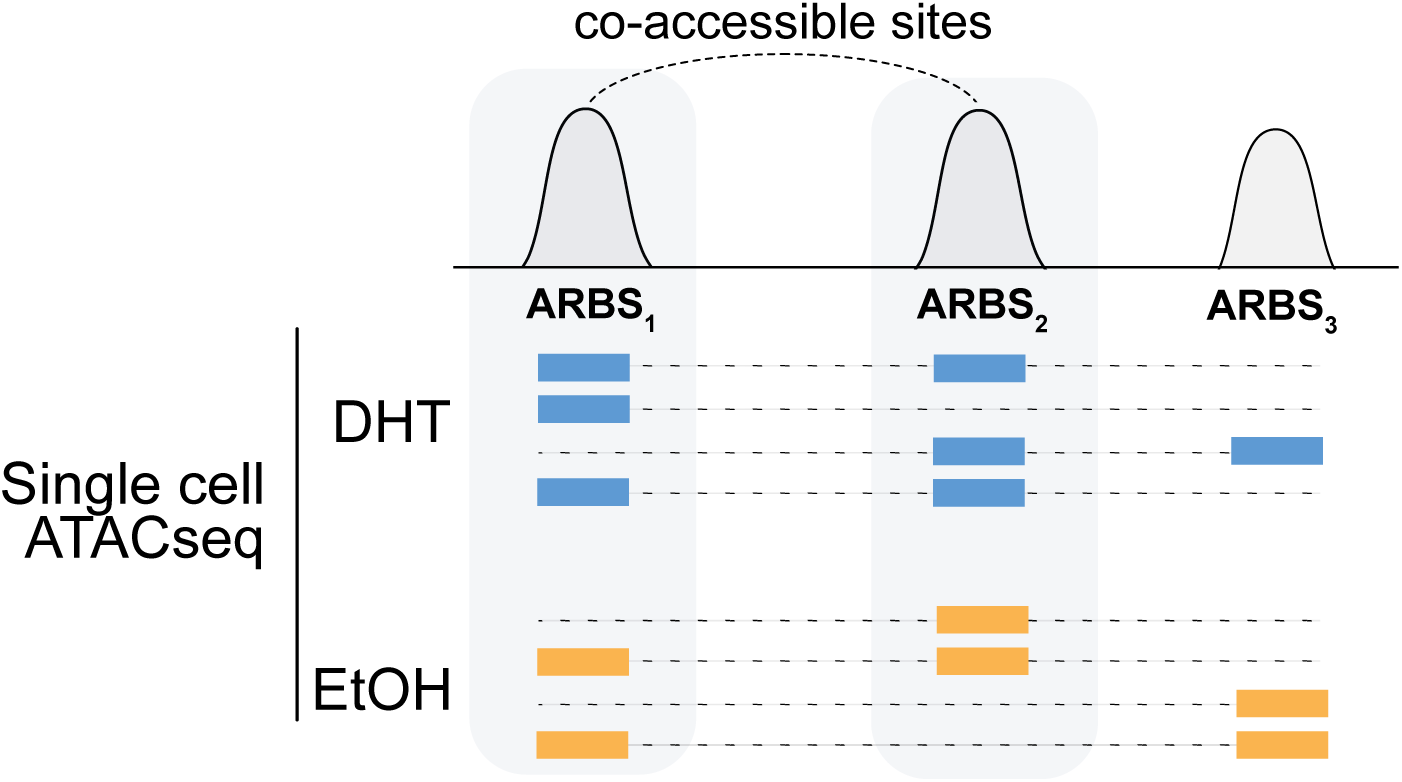
Schematic representation of co-accessibility quantification between ARBS using scATACseq data.

**Supplementary Figure 13:**
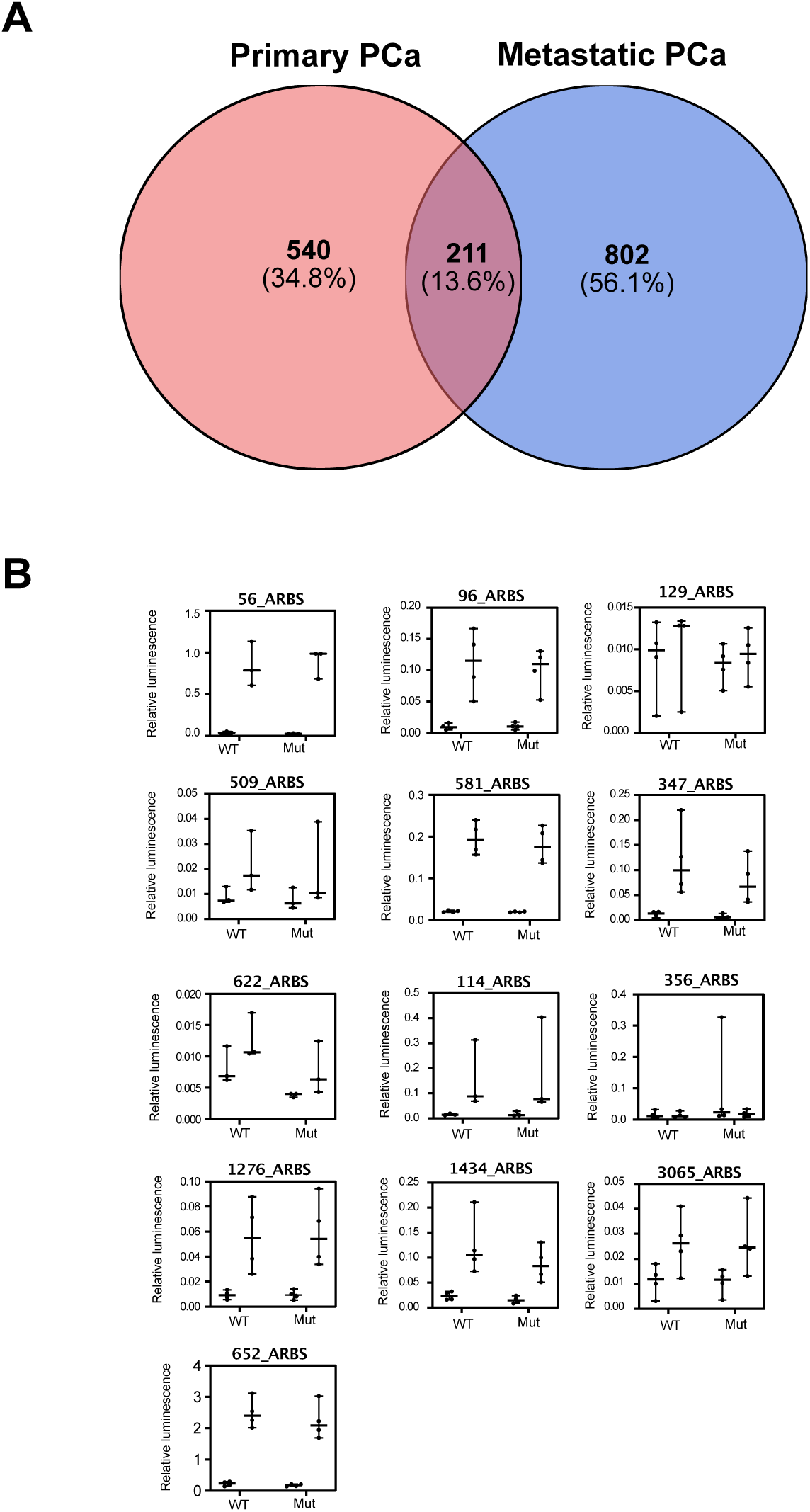
**(A)** Venn diagram of SNVs identified by WGS from primary PCa (n= 196) and metastatic CRPC (n=101). **(B)** Effects of SNV on the enhancer activity was quantified by the luciferase assay. From this 3/16 SNVs significantly affected androgen-mediated enhancer activity (Figure 6B). The remaining 13/16 SNVs did not significantly impact enhancer activity (*p-value>0.05,* 3 biological replica±SD).

