## Supplementary material for "Functional mapping of androgen receptor enhancer activity": https://docs.google.com/document/d/1w6qPf-SBIF2m9dmDQkOKcDi1sZAj6AMut8emfuFr7hE/edit

### Supplementary Figure 1

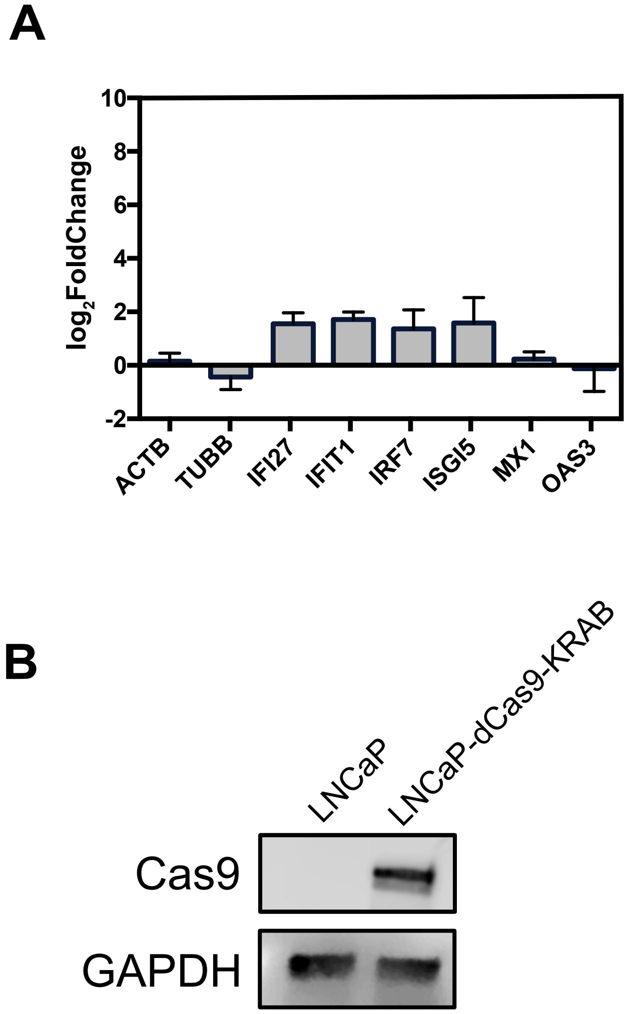

### Supplementary Figure 2

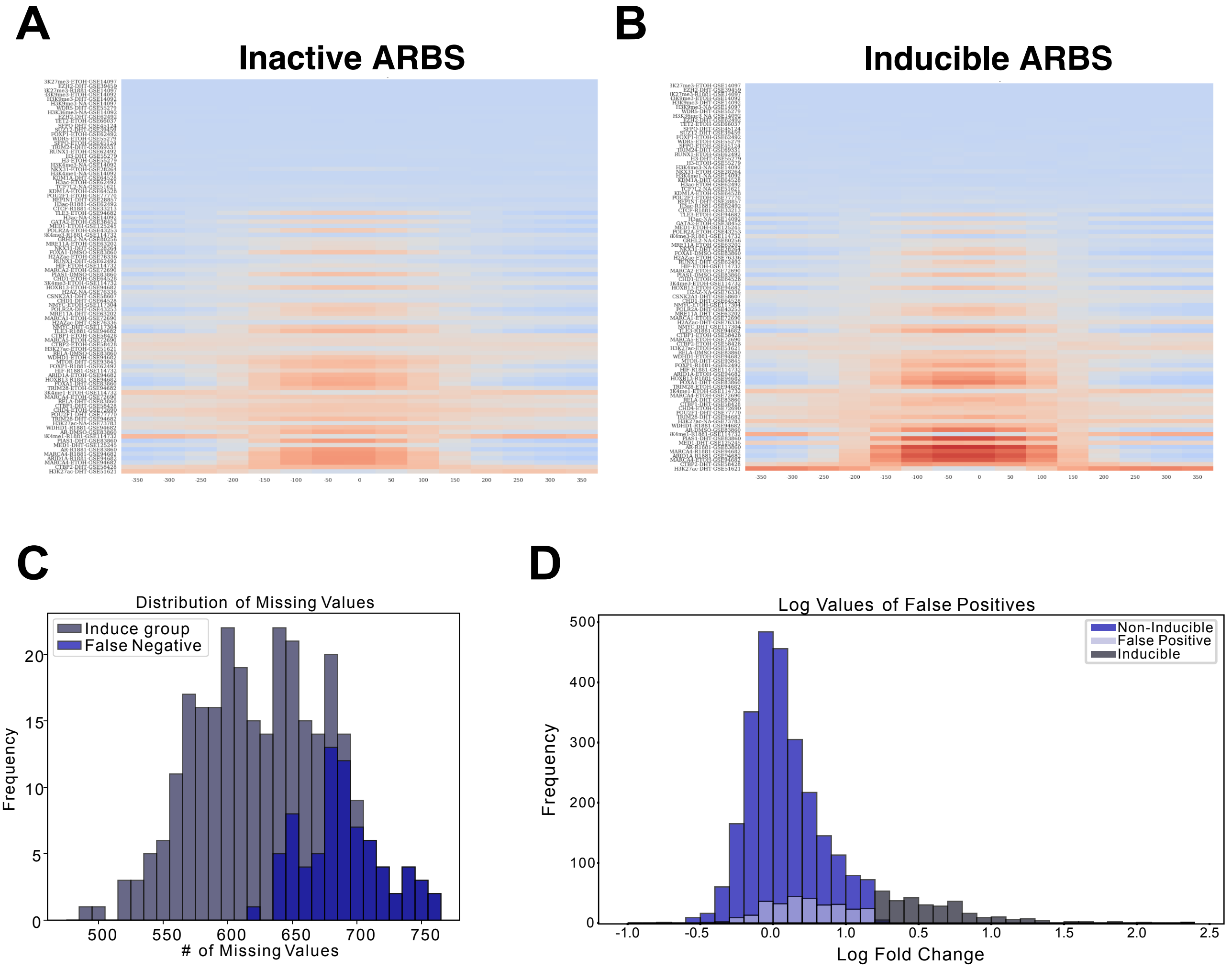

### Supplementary Figure 3

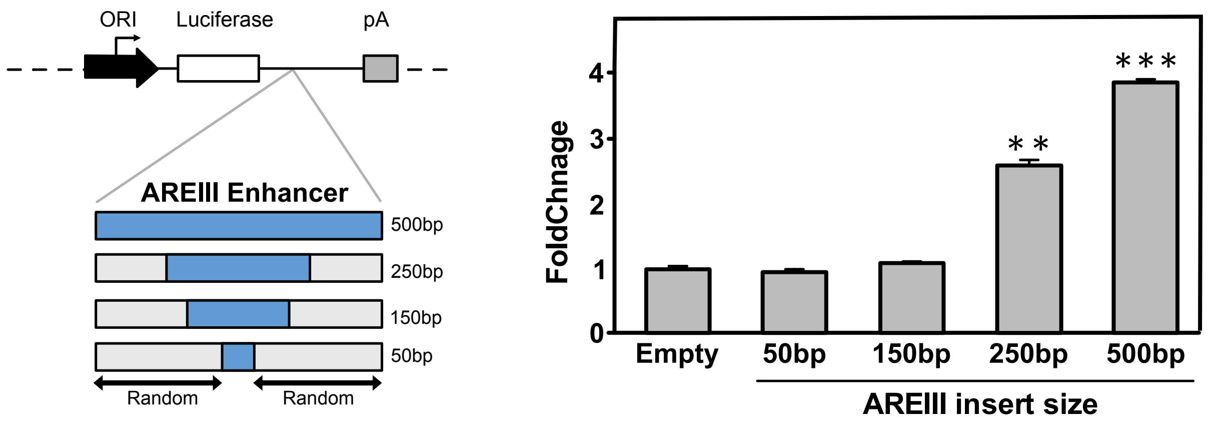

### Supplementary Figure 4

**A**

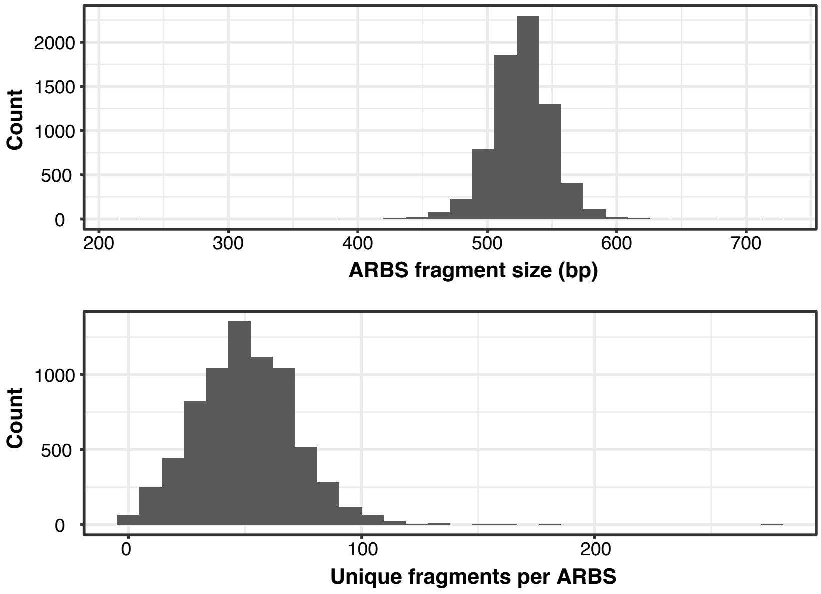


**B**

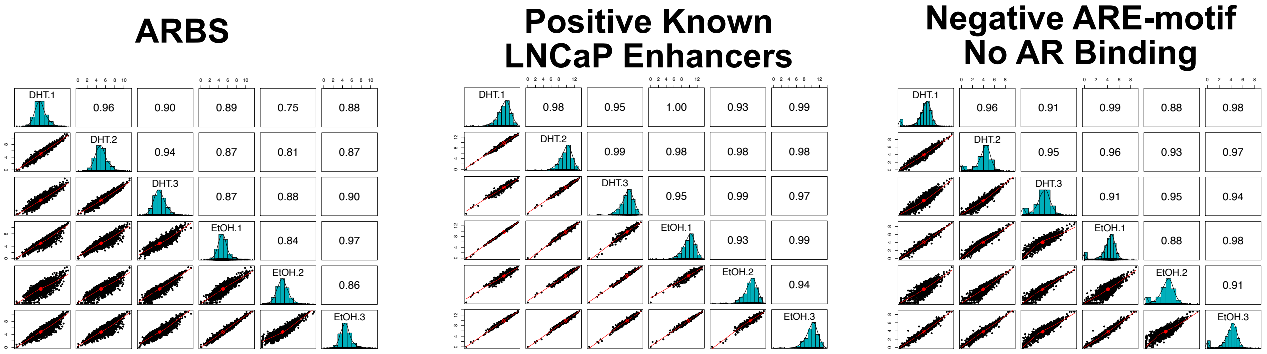

### Supplementary Figure 5

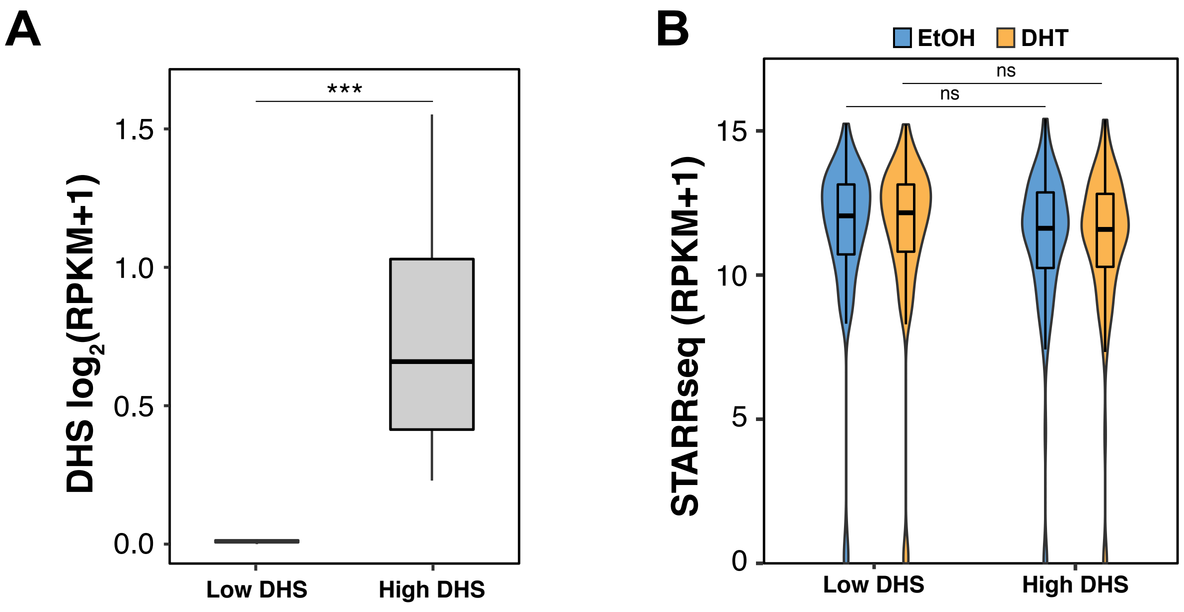

### Supplementary Figure 6

A

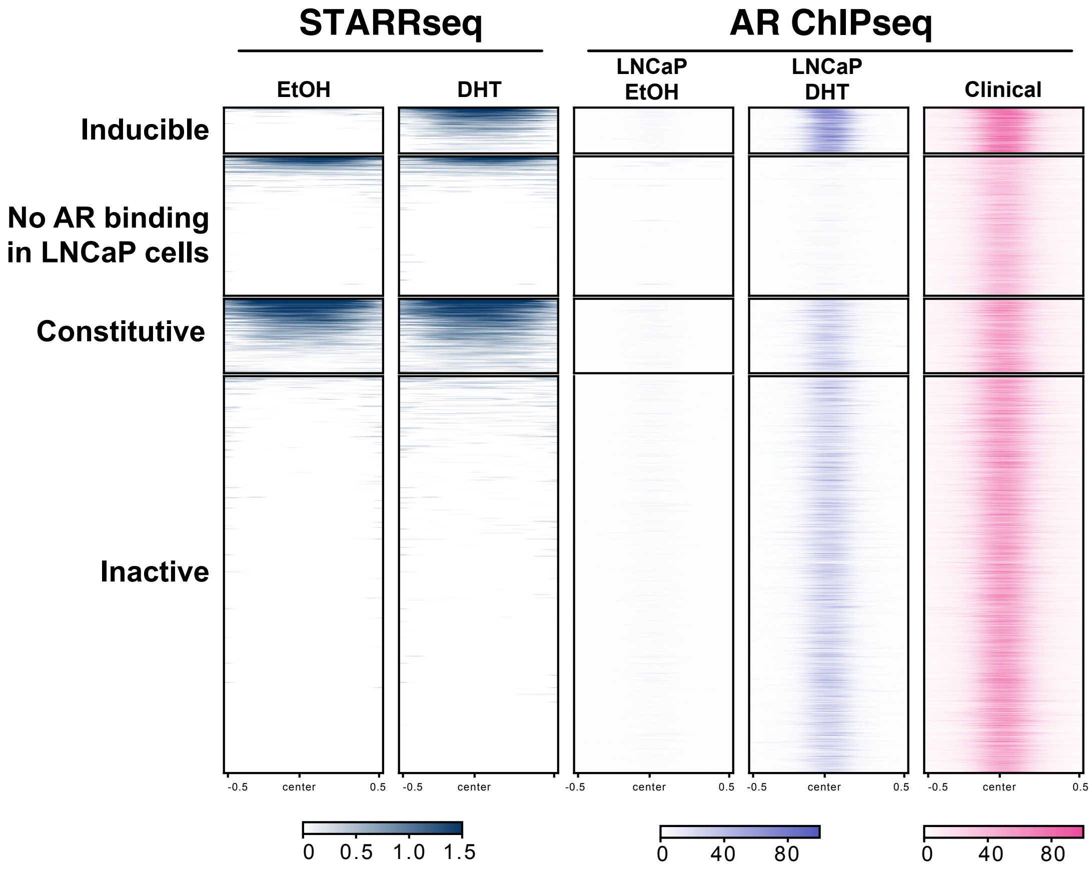


B

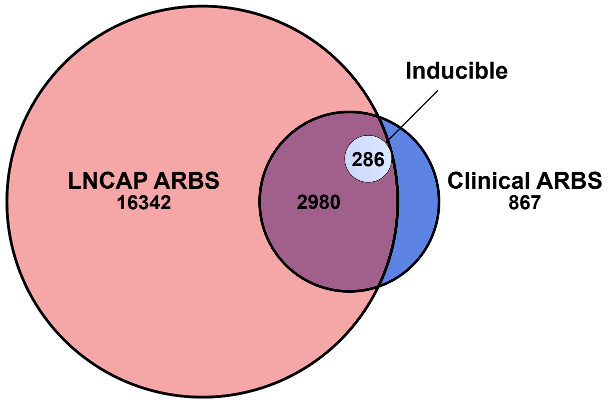

### Supplementary Figure 7

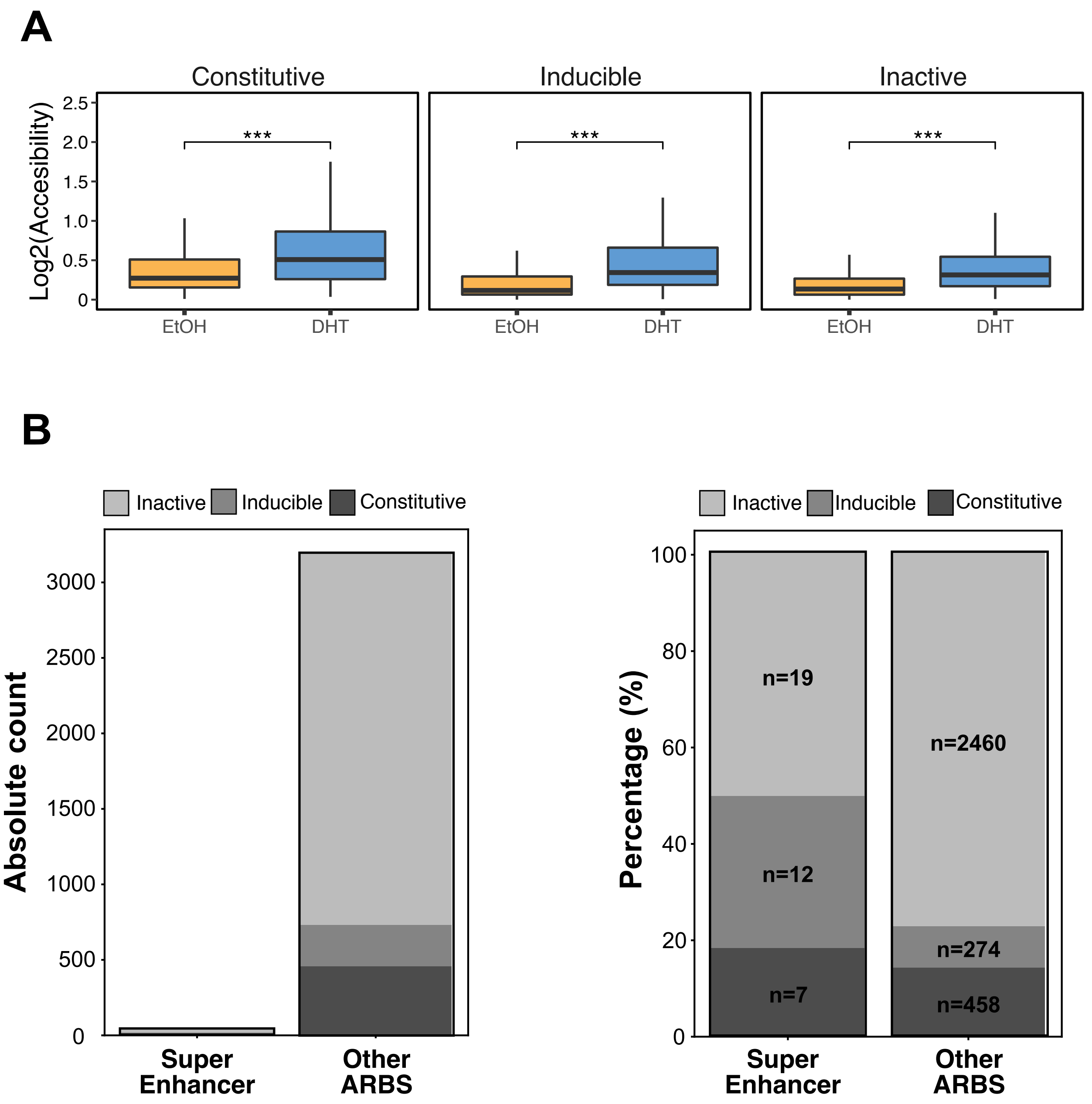

### Supplementary Figure 8

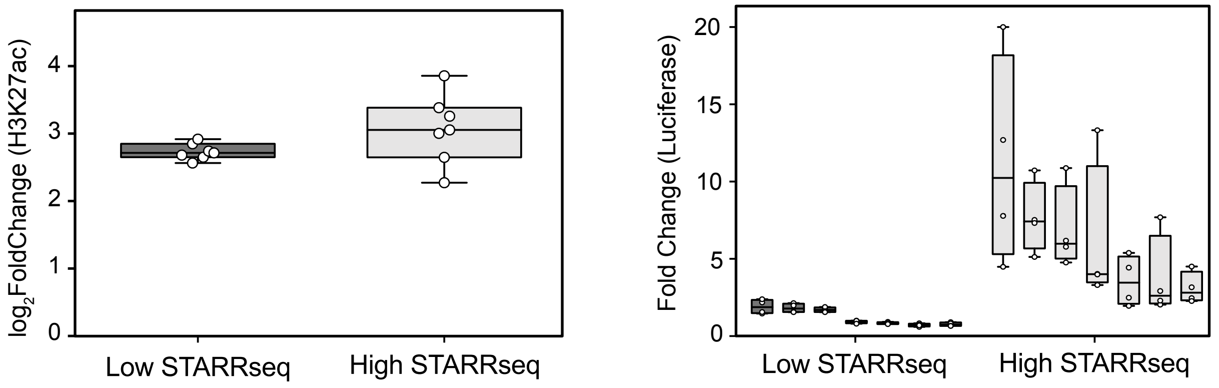

### Supplementary Figure 9

A

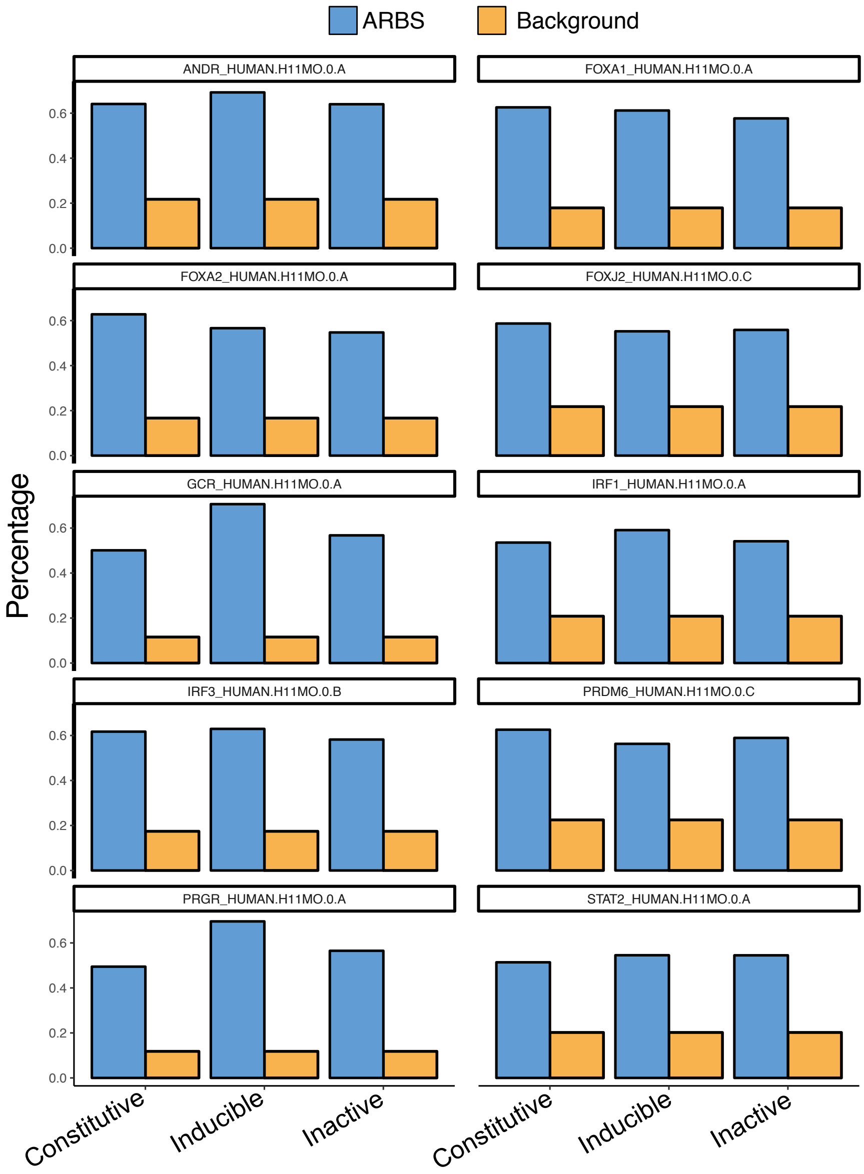


B

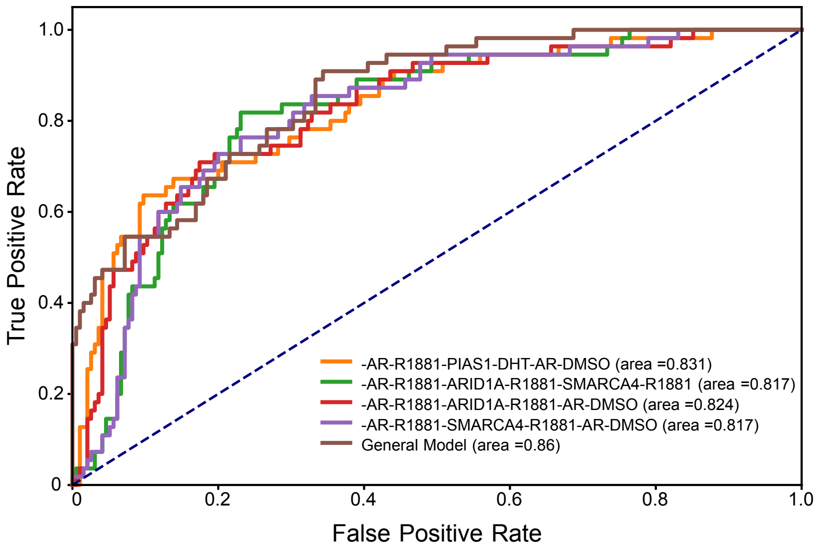

### Supplementary Figure 10

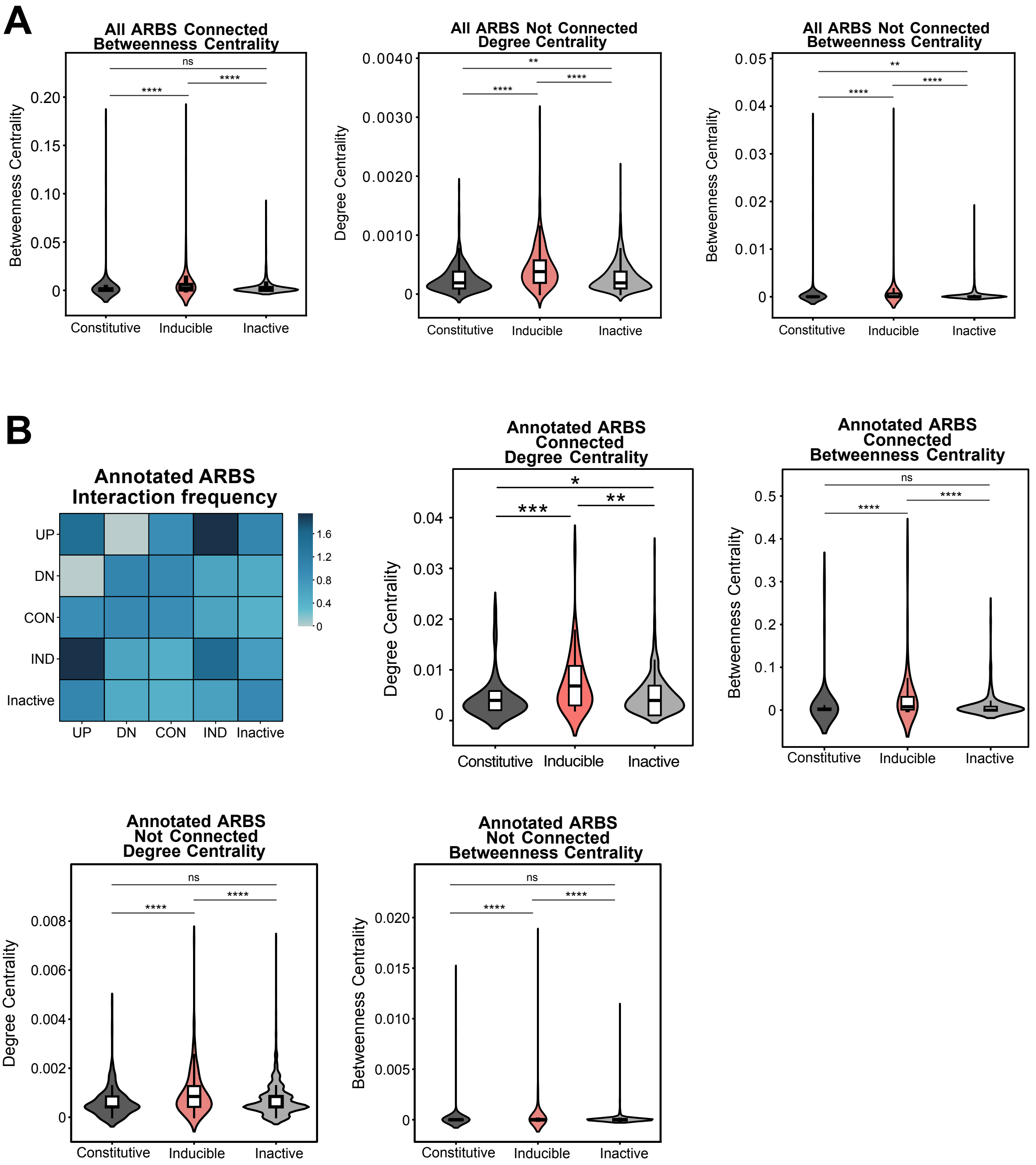

### Supplementary Figure 11

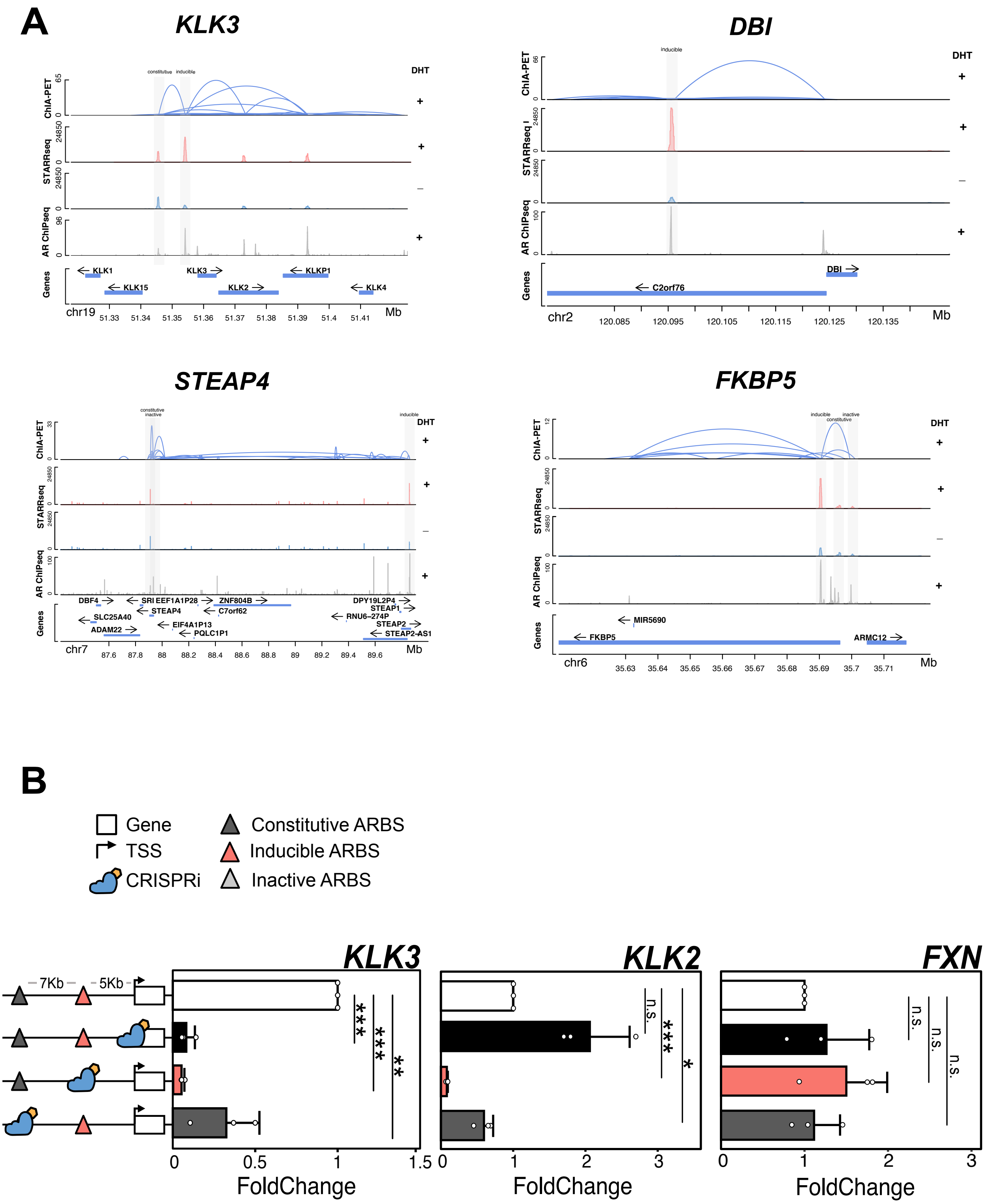

### Supplementary Figure 12

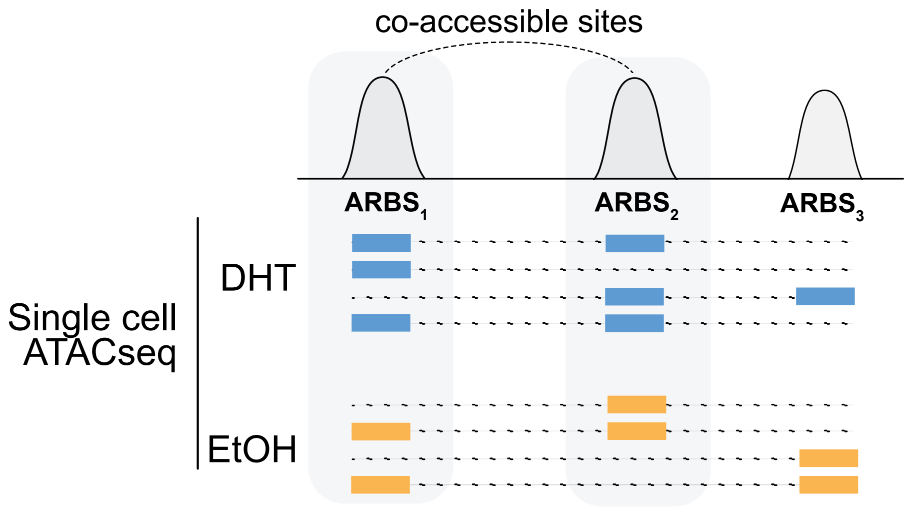

### Supplementary Figure 13

A

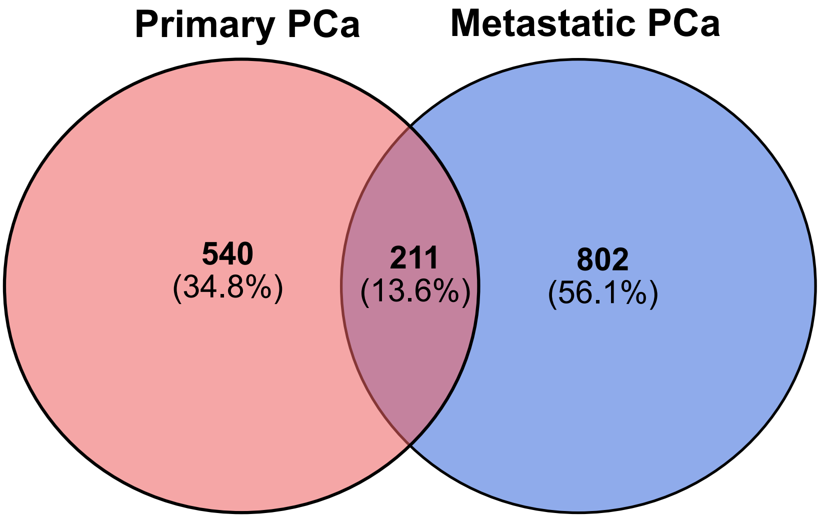


B

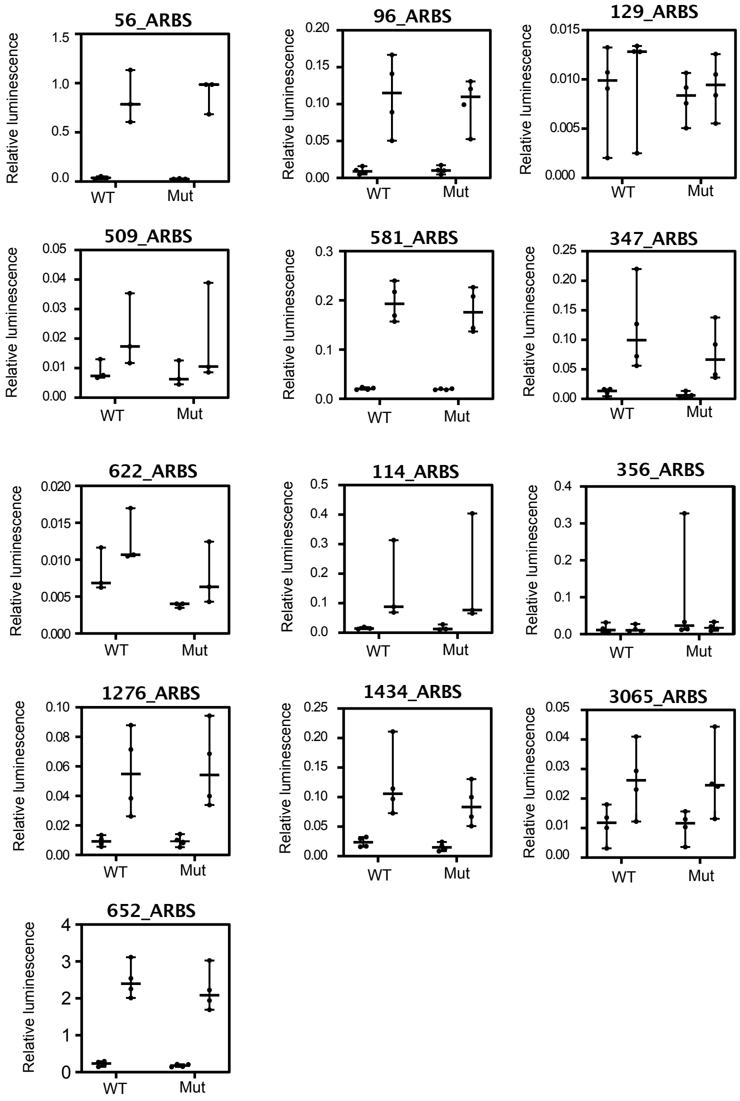
